# In-depth investigation of microRNA-mediated cross-kingdom regulation between Asian honey bee and microsporidian

**DOI:** 10.1101/2022.07.03.498592

**Authors:** Xiaoxue Fan, Wende Zhang, Kaiyao Zhang, Jiaxin Zhang, Qi Long, Ying Wu, Kuihao Zhang, Leran Zhu, Dafu Chen, Rui Guo

## Abstract

Asian honey bee *Apis cerana* is the original host for *Nosema ceranae*, a unicellular fungal parasite that causes bee nosemosis throughout the world. Currently, interaction between *A. cerana* and *N. ceranae* is largely unknown. Here, based on our previously gained high-quality RNA-seq and small RNA-seq data from *N. ceranae*-infected *A. c. cerana* workers’ midguts and clean spores, differentially expressed mRNAs (DEmiRNAs) in *N. ceranae* targeted by DEmiRNAs in host midguts and *A. c. cerana* DEmRNAs targeted by microsporidian DEmiRNAs were predicted using bioinformatics, and then target DEmRNAs in microsporidian and host were annotated and investigated, with a focus on targets involved in *N. ceranae* glycolysis/glyconeogenesis and virulence factors as well as *A. c. cerana* energy mechanism and immune response. It’s found that 97 down-regulated (60 up-regulated) mRNAs in NcCKM vs NcTM1 were potentially targeted by eight up-regulated (six down-regulated) miRNAs in AcCKMI1 vs AcTMI1, 44 down-regulated (15 up-regulated) mRNAs in NcCKM vs NcTM2 were putative targets of seven up-regulated (two down-regulated) miRNAs in AcCKMI2 vs AcTMI2. Additionally, miR-60-y and miR-676-y were found to up-regulate in AcCKMI1 vs AcTMI1 and target genes engaged in spore wall protein and glycolysis/gluconeogenesis, while miR-60-y in AcCKMI1 vs AcTMI1 was up-regulated and potentially targeted a glycolysis/gluconeogenesis-associated gene. Comparatively, 343 down-regulated (138 up-regulated) mRNAs in AcCKM1 vs AcTM1 were putative targets of 121 up-regulated (112 down-regulated) miRNAs in NcCKMI vs NcTMI1, 247 down-regulated (110 up-regulated) mRNAs were putatively targeted by 110 up-regulated (104 down-regulated) miRNAs in NcCKMI vs NcTMI2. Further analysis showed that 31 up-regulated miRNAs in NcCKMI vs NcTMI1 potentially targeted 12 down-regulated mRNAs in AcCKM1 vs AcTM1, which were involved in five immune-related pathways such as phagasome and Jak-STAT signaling pathway, whereas nine up-regulated miRNAs in NcCKMI vs NcTMI2 putatively targeted five down-regulated mRNAs in AcCKM2 vs AcTM2, which were engaged in three immune-related pathways including endocytosis, lysosomes, and regulation of autophagy. In addition, miR-21-x was observed to up-regulate in NcCKMI vs NcTMI1 and target a oxidative phosphorylation-related gene. Finally, potential targeting relationship between two host DEmiRNAs-microsporidian DEmRNAs pairs and two microsporidian DEmiRNAs-host DEmRNAs pairs were verified on basis of RT-qPCR. Our findings not only lay a foundation for exploring the molecular mechanism underlying cross-kingdom regulation between *A. c. cerana* workers and *N. ceranae*, but also offer valuable insights into Asian honey bee-microsporidian interaction.

## Introduction

*Apis cerana cerana*, a subspecies of the Asian honey bee, *Apis cerana*, is widely reared in China and many other Asian countries (Wu et al., 2020). Compared with western honey bee, *Apis mellifera, A. cerana* has several advantages such as adaptation to extreme weather conditions, collection of sporadic nectar sources, strong cleaning capability and colony-level defense ability, it’s therefore of special economic and ecological value (Arias and Sheppard, 2005; Gallai et al., 2009; Hepburn and Radloff, 2011). The reference genome of *A. cerana* had been published in 2015 (Pelin et al., 2015), laying a key foundation for further investigation of its biology and the behind molecular mechanisms (Park et al., 2015).

*Nosema ceranae* is an intracellular fungal parasite that exclusively infects the midgut epithelial cells of adult bees (Traver and Fell, 2011; Traver and Fell, 2012). Recent evidence suggested that *N. ceranae* was also infective for *A. mellifera* larvae (Eiri et al., 2015). *N. ceranae* can not only cause damage for host midgut cell structure, energy stress, immunosuppression, and inhibition of cell apoptosis, but also influence bee health and colony productivity in combination with other biotic or abiotic stress (Paris et al., 2018). The transmission of *N. ceranae* among individuals is mainly through faeces-oral or oral-oral route. A single coiled polar filament is highly compacted around the interior of the *N. ceranae* spore, and upon the stimulation by the environment within bee host midgut, the fungal spore germinates followed by ejection of the polar tube, which is then pieced into the host cell; the infective sporaplasm is transferred into the host cell and immediately starts the proliferation stage (Mayack et al., 2015; Martín-hernández et al., 2018). Similar to other microsporidia, after long-term evolution and co-adaptation, *N. ceranae* has evolved extremely reduced genome with a very small size and lost majority of pathways related to material and energy metabolism such as TCA circle and oxidative phosphorylation. Hence, *N. ceranae* is highly dependent on the host cell to provide the material and energy needed for fungal proliferation (Paris et al., 2018). Increasing evidence indicate that *N. ceranae* is able to enhance the synthesis of amino acids, lipids, and nucleotides during the infection process by releasing synthetic hexokinase into host cells, and at the meantime inhibits the host cell apoptosis to benefit their proliferation (Cuomo et al., 2012).

An array of transcriptomic studies had been performed to investigate response of *A. mellifera* workers to *N. ceranae* invasion. For example, Fu et al. (2019) performed transcriptome investigation of immune response of *A. m. ligustica* workers to *N. ceranae* infection, and revealed that genes encoding apideacin, defensin-1, and hymenoptaecin were differentially expressed during the infection process. However, few omics studies were focused on *N. ceranae* during the infection process. To clarify the mechanisms of *N. ceranae* parasitism, Huang et al. (2016b) conducted deep sequencing and time-series analysis of *N. ceranae*-infected *A. mellifera* workers’ midguts, the result indicated that 1 122 microsporidian genes clustered into four expression patterns were significantly differently expressed during the fungal reproduction cycle. Based on dissection of transcriptomic dynamics of *N. ceranae* infecting *A. m. ligustica* workers, our group uncovered that genes encoding virulence factors such as spore wall protein and ricin B lectin were likely to play crucial roles in microsporidian proliferation (Geng et al., 2020c).

MicroRNAs (miRNAs) are a kind of single-strand small non-coding RNAs (ncRNAs), with a length distribution among 19-25 nt. MiRNAs play a regulatory part in gene expression at post-transcriptional level, leading to mRNA cleavage or translational suppression (Garofalo et al., 2011). MiRNAs have been suggested to participate in a substantial quantity of biological processes such as cell differentiation and immune response (Bartel, 2004; Pillai et al., 2005). Recent documentations demonstrated that miRNAs could not only regulate the endogenous gene expression, but also modulate the expression of exogenous genes. Zhang et al. (2012) discovered that plant-derived MIR168a and MIR156a were stably expressed in human and mice and negatively-regulated the expression of the target gene encoding LDLRAP1. This was the first report of miRNA-mediated cross-kingdom regulation between plant and animal. Hereafter, increasing studies confirmed cross-kingdom regulation among animals, plants, and microorganisms (Zhang et al., 2016, Liu et al., 2012, Cui et al., 2019). However, study on cross-regulation between bees and pathogens is still very limited. After silencing *Dicer* gene in *N. ceranae* with specific siRNA, Evans and Huang (2018) performed next-generation sequencing and analysis of *A. mellifera* workers and *N. ceranae* across a full fungal proliferation cycle, the result suggested that *N. ceranae* miRNAs may regulate expression of genes in both parasite and host. Recently, on basis of transcriptome sequencing and bioinformatics, our team conducted comprehensive investigation of miRNA-mediated regulation between *A. m. ligustica* workers and *N. ceranae* during the infection process (Du et al., 2021; Fan et al., 2021).

Currently, miRNA-mediated cross-kingdom regulation between eastern honey bee and microsporidian is completely unknown. Here, based on previously gained high-quality transcriptome data, differentially expressed mRNAs (DEmRNAs) in microsporidian targeted by *A. c. cerana* DEmiRNAs and host DEmRNAs targeted by *N. ceranae* DEmiRNAs were for the first time predicted and analyzed, followed by in-depth investigation of DEmiRNA-mediated cross-kingdom regulation between host and microsporidian. Our results will not only lay a key foundation for clarifying the underlying molecular mechanism but also offer a new insight into Asian honey bee-microsporidian interaction.

## Materials and method

### Source of sRNA-seq and RNA-seq data from *A. c. cerana* workers’ midguts

In our previous work, midgut tissues of *N. ceranae*-inoculated *A. c. cerana* workers at 7 days post inoculation (dpi) and 10 dpi (AcTMI1 group: AcTMI1-1, AcTMI1-2, AcTMI1-3; AcTMI2 group: AcTMI2-1, AcTMI2-2, AcTMI2-3) and corresponding un-inoculated workers’ midgut tissues (AcCKMI1 group: AcCKMI1-1, AcCKMI1-2, AcCKMI1-3; AcCKMI2 group: AcCKMI2-1, AcCKMI2-2, AcCKMI2-3) were prepared, followed by construction of cDNA libraries and deep sequencing utilizing small RNA sequencing (sRNA-seq) technology. Totally, 122,104,443 clean reads were obtained after quality control of raw reads generated from sRNA-seq, and the Pearson correlation coefficient between every biological replica in each group was above 96.19% (Du et al., 2019). Raw data were deposited in the Sequence Read Archive (SRA) database (http://www.ncbi.nlm.nih.gov/sra/) and connected to BioProject: PRJNA487111. The high-quality sRNA-seq data can provide a basis for prediction and analysis of *N. ceranae* DEmRNAs targeted by *A. c. cerana* DEmiRNA in this study.

In another previous study, midgut tissues of *N. ceranae*-inoculated *A. c. cerana* workers at 7 days post inoculation (7 dpi) and 10 dpi (AcTM1 group: AcTM1-1, AcTM1-2, AcTM1-3; AcTM2 group: AcTM2-1, AcTM2-2, AcTM2-3) and corresponding un-inoculated workers’ midgut tissues (AcCKM1 group: AcCKM1-1, AcCKM1-2, AcCKM1-3; AcCKM2: AcCKM2-1, AcCKM2-2, AcCKM2-3) were prepared, followed by strand-specific cDNA library construction and deep sequencing using RNA-seq technology. Quality control result indicated that In total, 1 562 162 742 clean reads with an average Q30 of 94.76% were obtained after quality control of raw reads produced from RNA-seq; additionally, 121 949 977 (75.78%), 171 868 060 (55.01%), 97 432 266 (78.13%), and 200 570 776 (44.19%) clean reads in AcCKM1, AcTM1, AcCKM2, and AcTM2 groups were mapped to the reference genome of *A. cerana* (Assembly ACSNU-2.0), respectively; moreover, the Pearson correlation coefficient between every biological replica in each group was above 0.87 (Xing et al., 2021). The RNA-seq data with high quality could be used for prediction and investigation of *A. c. cerana* DEmRNAs targeted by *N. ceranae* DEmiRNAs.

### Sources of sRNA-seq and RNA-seq data from *N. ceranae* spores

*N. ceranae* spores (NcCKMI group: NcCKMI-1, NcCKMI-2, NcCKMI-3) were previously prepared with Percoll discontinuous density gradient centrifugation method by our group, followed by construction of cDNA libraries and deep sequencing using sRNA-seq(Geng et al., 2020a). Raw data from sRNA-seq were uploaded to the NCBI SRA database under BioProject numbers: PRJNA395264.

Meantime, the aforementioned *N. ceranae* spores (NcCKM group: NcCKM-1, NcCKM-2, NcCKM-3) were subjected to cDNA library construction and by RNA-seq. Following strand-specific cDNA library-based RNA-seq, over 218.47 million raw reads were generated from three cDNA libraries of *N. ceranae* spores after removing adaptor sequences and low-quality reads; additionally, 107 123 113, 117 688 499 and 116 776 007 clean reads were mapped to the *N. ceranae* reference genome (assembly ASM18298v1); Original data were uploaded to the NCBI SRA databased and linked to BioProject number: PRJNA395264 (Guo et al., 2018)

The high-quality sRNA-seq and RNA-seq data could be used for identification of DEmiRNAs and DEmRNAs in *N. ceranae* during the infection of *A. c. cerana* workers. and prediction and investigation of *N. ceranae* DEmRNAs targeted by *A. c. cerana* DEmiRNAs as well as *N. ceranae* DEmiRNA-targeted *A. c. cerana* DEmRNAs.

### Source of sRNA-seq and RNA-seq data from *N. ceranae* infecting *A. c. cerana* workers

On basis of our established protocol (Du et al., 2019), clean tags from *N. ceranae* infecting *A. c. cerana* workers at 7 dpi and 10 dpi (NcTMI1 group: NcTMI1-1, NcTMI1-2, NcTMI1-3; NcTMI2 group: NcTMI2-1, NcTMI2-2, NcTMI2-3) were filtered out from sRNA-seq data of midgut tissues of *A. c. cerana* workers at 7 dpi and 10 dpi. Briefly, (1) the clean tags from sRNA-seq of midgut tissues of *A. c. cerana* workers at 7 dpi and 10 dpi were first mapped to the GeneBank and Rfam databases to remove ribosomal RNA (rRNA), small cytoplasmic RNA (scRNA), small nucleolar RNA (snoRNA), small intranuclear RNA (snRNA), and transport RNA (tRNA) data; (2) the unmapped clean reads were then mapped to the *A. cerana* reference genome (Assembly ACSNU-2.0) by using Bowtie software (Langmead et al., 2009) to remove host-derived data; (3) the unmapped clean tags were further mapped to the *N. ceranae* reference genome (Assembly ASM98816v1), and the mapped data were derived from *N. ceranae* during the infection. In total, 36 792 918 and 39 888 369 clean tags from NcTMI1, and NcTMI2 groups, respectively. Original data were uploaded to the NCBI SRA databased and linked to BioProject number: PRJNA406998.

Following our previous described method, clean reads from *N. ceranae* infecting *A. c. cerana* workers at 7 dpi and 10 dpi (NcTM1 group: NcTM1-1, NcTM1-2, NcTM1-3; NcTM2 group: NcTM2-1, NcTM2-2, NcTM2-3) were filtered out from RNA-seq data of midgut tissues of *A. c. cerana* workers at 7 dpi and 10 dpi. Totally, 1 562 162 742 clean tags derived from *N. ceranae* during the infection with mean Q30 of 92.63% were gained (Xiong et al., 2020).Original data were uploaded to the NCBI SRA databased and linked toBioProject number: PRJNA562784

### Prediction and analysis of *N. ceranae* DEmRNAs targeted by *A. c. cerana* DEmiRNA

The expression level of each *A. c. cerana* miRNA was normalized to the total number of sequence tags per million (TPM) following the formula: normalized expression = mapped read count/total reads × 10^6^. edgeR software (Robinson et al., 2010) was used to screen out the DEmiRNAs in AcCKMI1 vs AcTMI1 and AcCKMI2 vs AcTMI2 comparison groups following the criteria of *P* value (FDR corrected) < 0.05 and |log_2_(Fold change)| > 1.

The expression level in each *N. ceranae* mRNA was normalized to the fragments per kilobase of transcript per million fragments mapped (FPKM) following the formula: total exon Fragments / (mapped reads(Millions) × exon length(KB)). DEmRNAs in NcCKM vs NcTM1 and NcCKM vs NcTM2 comparison groups were screened out based on the criteria of *P* value (FDR corrected) < 0.05 and |log_2_(Fold change)| > 1.

*N. ceranae* DEmRNAs targeted by *A. c. cerana* DEmiRNAs were predicted using TargetFinder software (Allen et al., 2005) with default parameters. Gene Ontology (GO) classification and Kyoto Encyclopedia of Genes and Genomes (KEGG) pathway analysis of the aforementioned *A. c. cerana* DEmiRNAs was performed using related tools in OmicShare platform (http://www.omicshare.com/tools/). Regulatory networks between *A. c. cerana* DEmiRNAs and *N. ceranae* DEmRNAs constructed based on targeting relationship followed by visualization using Cytoscape v.3.2.1 software (Smoot et al, 2011) with default parameters.

### Prediction and investigation of *A. c. cerana* DEmRNAs targeted by *N. ceranae* DEmiRNAs

DEmiRNAs in NcCKMI vs NcTMI1 and NcCKMI vs NcTMI2 comparison groups and DEmRNAs in NcCKM1 vs NcTM1 and NcCKM2 vs NcTM2 comparison groups were identified following the above-mentioned methods.

*A. c. cerana* DEmRNAs targeted by *N. ceranae* DEmiRNAs were predicted with TargetFinder software. GO categorization and KEGG pathway analysis of *A. c. cerana* DEmRNAs were conducted using OmicShare platform. Regulatory networks between *N. ceranae* DEmiRNAs and *A. c. cerana* DEmRNAs were constructed on basis of the targeting relationship and then visualized with Cytoscape v.3.2.1 software.

### RT-qPCR Validation of DEmRNAs

To verify the reliability of the transcriptome datasets used in this study, according to the targeted binding relationship, two host DEmRNAs (XM_017055873.1, XM_017064721.1) in AcCKM1 vs AcTM1, two microsporidian DEmiRNAs (miR-8462-x, miR-676-y) in NcCKMI vs NcTMI1, two microsporidian DEmRNAs (XM_002995668.1, XM_002995068.1) in NcCKM vs NcTM1 and two host DEmiRNAs in AcCKM1 vs AcTM1 were randomly selected for RT-qPCR. The first cDNA strand was synthesized with the SuperScript first-strand synthesis system (Yeasen) according to the protocol. Primers for qPCR were designed utilizing DNAMAN software and synthesized by Sangon Biotech Co., Ltd. (Shanghai). The housekeeping gene actin was used as an internal control. The RNA samples used as templates for RNA-seq were the same as those used for RT-qPCR, which was conducted on a QuanStudio Real-Time PCR System (ThemoFisher, Walthem, MA, USA). The 20 μL PCR reaction mixture contained 10 μL SYBR Green dye (Yeasen), 1 μL (10 μmol/L) specific forward primer, 1 μL (10 μmol/L) reverse primer, 1 μL (10 ng/μL) diluted cDNA, and 7 μL RNase free water. Cycling parameters were as follows: 95 °C for 1 min, followed by 40 cycles at 95 °C for 15 s, 55 °C for 30 s, and 72 °C for 45 s. The relative gene expression was calculated using the 2^-ΔΔCT^ method. The experiment was carried out times using three independent biological samples.

## Result

### Analysis of *N. ceranae* DEmRNAs targeted by DEmiRNAs in *A. c. cerana* workers’ midguts and corresponding regulatory networks

Target prediction suggested that 97 down-regulated mRNAs in NcCKM vs NcTM1 were potentially targeted by eight up-regulated miRNAs in AcCKMI1 vs AcTMI1 (**Figure 1A, see also Table S1**), whereas 60 up-regulated *N. ceranae* mRNAs were potential targets of six down-regulated *A. c. cerana* miRNAs (**Figure 1B, see also Table S1**). The aforementioned 97 down-regulated mRNAs were related to eight biological process-associated terms including metabolic process and cellular process, six cellular component-associated terms including cell and cell part, two molecular function-associated terms including binding and catalytic activity (**Table S3**); these down-regulated mRNAs were also annotated to 35 pathways including metabolic pathways, biosynthesis of antibiotics, and biosynthesis of secondary metabolites (**Table S4**). Additionally, the 60 up-regulated mRNAs were associated with 16 functional terms such as metabolic process, cell part, and catalytic activity (**Table S3**); and 33 pathways such as metabolic pathway, biosynthesis of antibiotics, and biosynthesis of secondary metabolites (**Table S4**).

**FIGURE 1.**
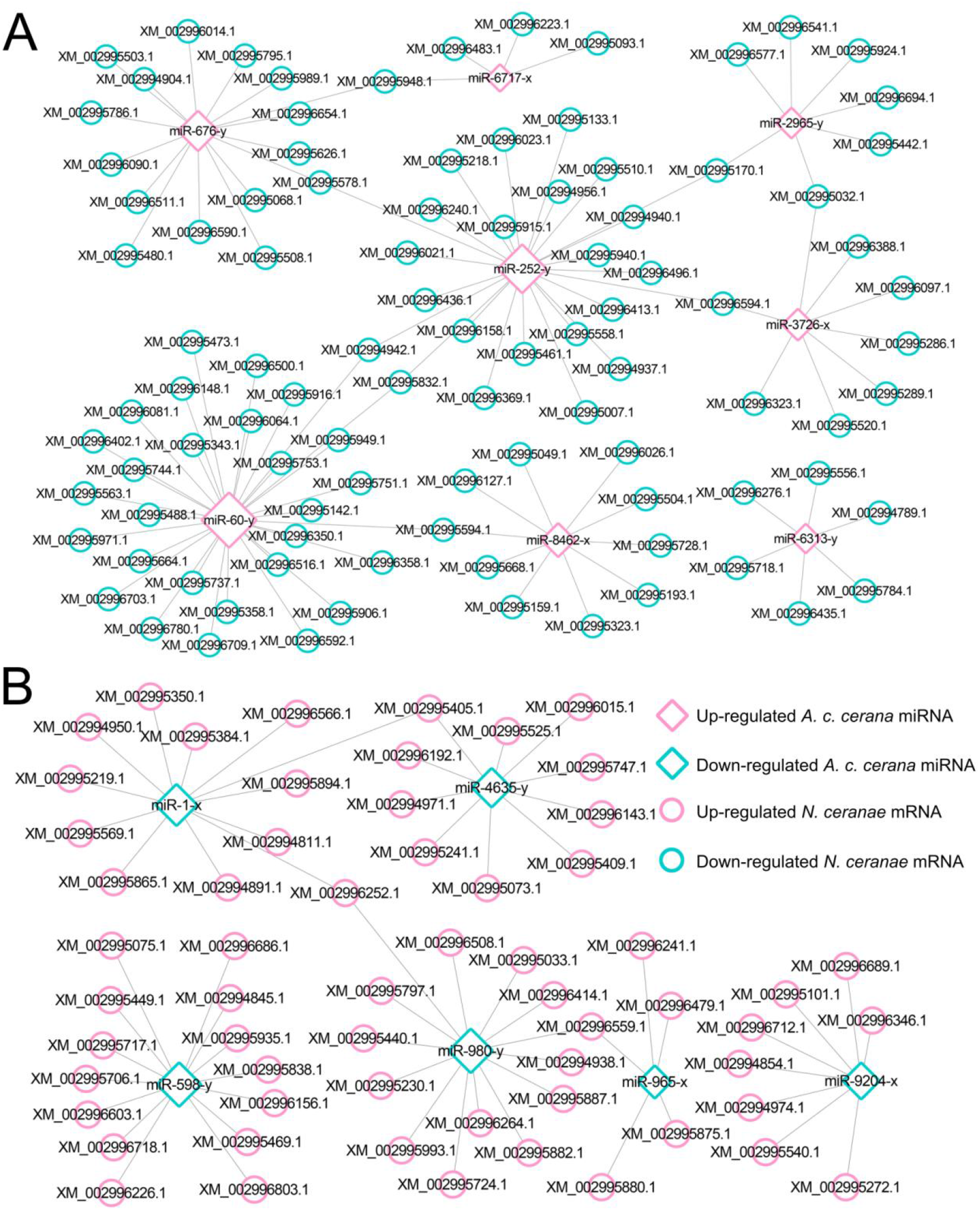
Regulatory network between *A. c. cerana* DEmiRNAs and target DEmRNAs in *N. ceranae* at 7 dpi. **(A)** Regulatory network between up-regulated miRNAs in host and down-regulated mRNAs in microsporidian. **(B)** Regulatory network between down-regulated miRNAs in host and up-regulated mRNAs in microsporidian.

In NcCKM vs NcTM2 comparison group, 44 down- and 15 up-regulated mRNAs were potentially targeted by seven up- and two down-regulated miRNAs in AcCKMI2 vs AcTMI2 comparison group (**Figure 2, see also Table S1**). The above-mentioned 15 up-regulated *N. ceranae* mRNAs were relative to eight functional terms including catalytic activity, binding, and metabolic process (**Tabel S3**); and 11 pathways such as metabolic pathways, biosynthesis of secondary metabolites and carbon metabolism (**Tabel S4**). Additionally, the 44 down-regulated *N. ceranae* mRNAs were engaged in eight functional terms such as metabolic process, cellular process, and single-organism process(**Table S3**); and 24 pathways including metabolic pathways, ribosome biogenesis in eukaryotes, and endocytosis(**Table S4**).

**FIGURE 2.**
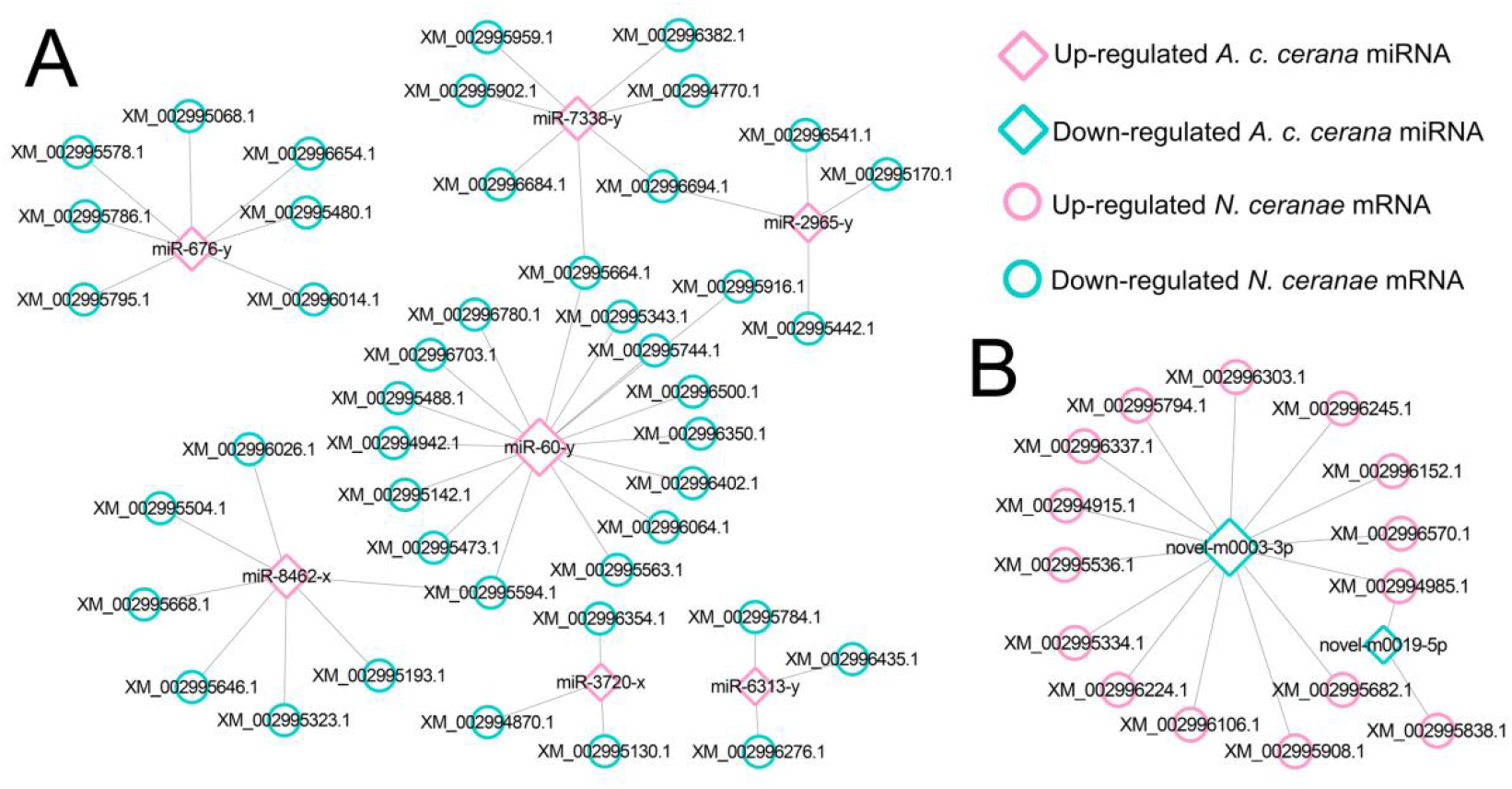
Regulatory network between *A. c. cerana* DEmiRNAs and target DEmRNAs in N. cerana at 10 dpi. **(A)** Regulatory network between up-regulated miRNAs in host and down-regulated mRNAs in microsporidian. **(B)** Regulatory network between down-regulated miRNAs in host and up-regulated mRNAs in microsporidian.

Further investigation indicated that five up-regulated miRNAs shared by AcCKMI1 vs AcTMI1 and AcCKMI2 vs AcTMI2 comparison groups, which potentially targeted 35 down-regulated mRNAs shared by NcCKM vsNcTM1 and NcCKM vsNcTM2 comparison groups (**Figure 3**).

**FIGURE 3.**
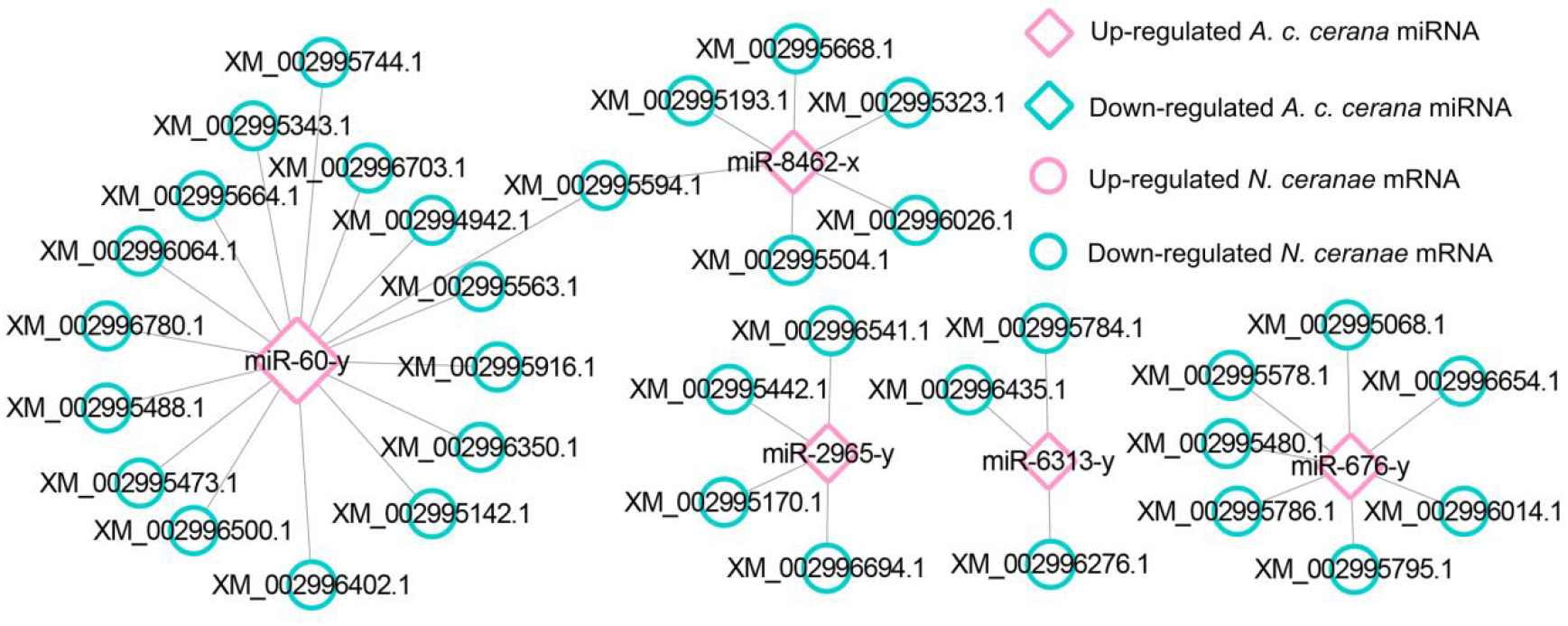
Regulatory network between up-regulated miRNAs shared by AcCKMI1 vs AcTMI1 and AcCKMI2 vs AcTMI2 and down-regulated mRNAs shared by NcCKM vs NcTM1 and NcCKM vs NcTM2.

The aforementioned common down-regulated *N. ceranae* mRNAs were annotated to 14 GO categories, including seven biological process-associated categories such as cellular process and metabolic process, five cellular component-associated categories such as cell and cell part, two molecular function-associated categories such as catalytic activity and binding (**Figure 4A**). In addition, these common down-regulated mRNAs were also annotated 20 pathways, among which the most abundant one was metabolic pathway followed by ribosome biogenesis in eukaryotes and pyrimidine metabolism (**Figure 4B**).

**FIGURE 4.**
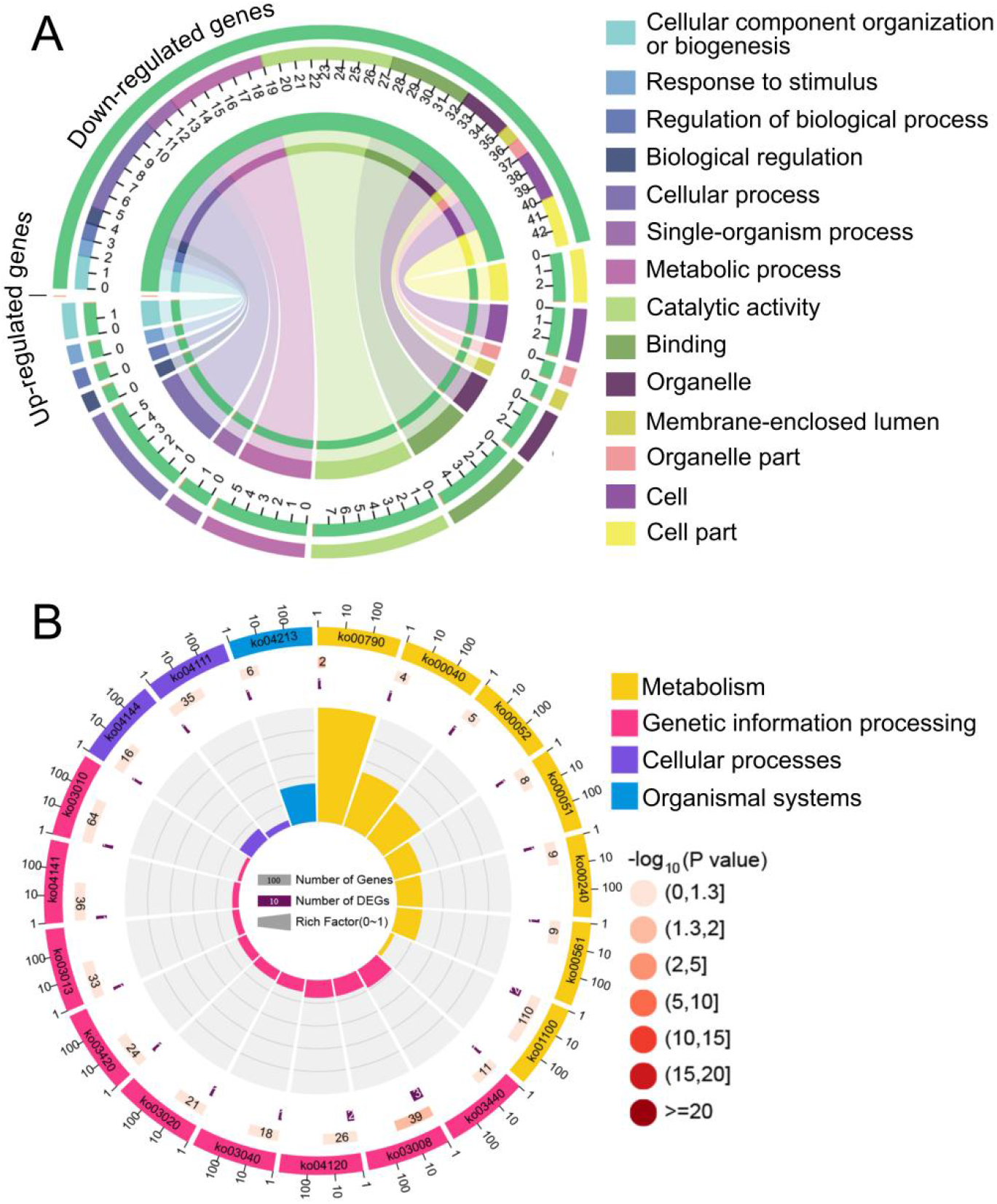
Database annotation of shared down-regulated *N. ceranae* mRNAs targeted by shared up-regulated *A. c. cerana* miRNAs. **(A)** GO database annotation of down-regulated mRNAs shared by NcCK1 vs NcTM2 and NcCK1 vs NcTM2. **(B)** KEGG database annotation of down-regulated mRNAs shared by NcCK1 vs NcTM2 and NcCK1 vs NcTM2.

### Investigation of *A. c. cerana* DEmiRNAs and their target DEmRNAs associated with *N. ceranae* virulence factors

In AcCKMI1 vs AcTMI1 comparison group, the up-regulated miR-60-y (log_2_FC=10.87, *P*<0.01) potentially targeted a spore wall protein-encoding gene (XM_002996592.1, log_2_FC=-2.24, *P*<0.01) in NcCK1 vs NcTM1 comparison group; the up-regulated miR-676-y (log_2_FC=12.97, *P*<0.01) potentially targeted a gene encoding pyruvate dehydrogenase e1 component subunit alpha (XM_002996090.1, log_2_FC=-6.89, *P*<0.01) relative to glycolysis/gluconeogenesis pathway. In AcCKMI2 vs AcTMI2 comparison group, the miR-60-y (log_2_FC=14.32, *P*<0.01) potentially targeted an acs1p-encoding gene associated with glycolysis/gluconeogenesis pathway (XM_002995904.1, log_2_FC=3.78, *P*<0.01) in NcCK2 vs NcTM2 comparison group.

### Analysis of DEmRNAs in *A. c. cerana* workers’ midguts targeted by *N. ceranae* DEmiRNAs

In total, 343 down-regulated mRNAs in AcCKM1 vs AcTM1 were putative targets of 121 up-regulated miRNAs in NcCKMI vs NcTMI1 (**Figures 5A, see also Table S2**), while 138 up-regulated mRNAs of *A. c. cerana* in AcCKM1 vs AcTM1 were potentially targeted by 112 down-regulated in NcCKMI vs NcTMI1 (**Figures 5B, see also Table S2**). The aforementioned 343 down-regulated mRNAs were related to 15 biological process-associated functional terms including cellular process and metabolic process, ten cellular component-associated functional terms including cell and cell part, six molecular function-associated functional terms including binding and catalytic activity (**Tabel S3**); and 217 pathways such as Jak-STAT signaling pathway, endocytosis, and lysosome (**Tabel S4**). Additionally, 138 up-regulated mRNAs were associated with 11 biological process-related functional terms such as cellular process, single-organism process, and metabolic process, eight cellular component-associated functional terms including membrane part, membrane and organelle, seven molecular function-associated functional terms including binding, catalytic activity and molecular transducer activity (**Tabel S3**); and 107 pathways such as metabolic pathways, oxidative phosphorylation, and purine metabolism (**Tabel S4**).

**FIGURE 5.**
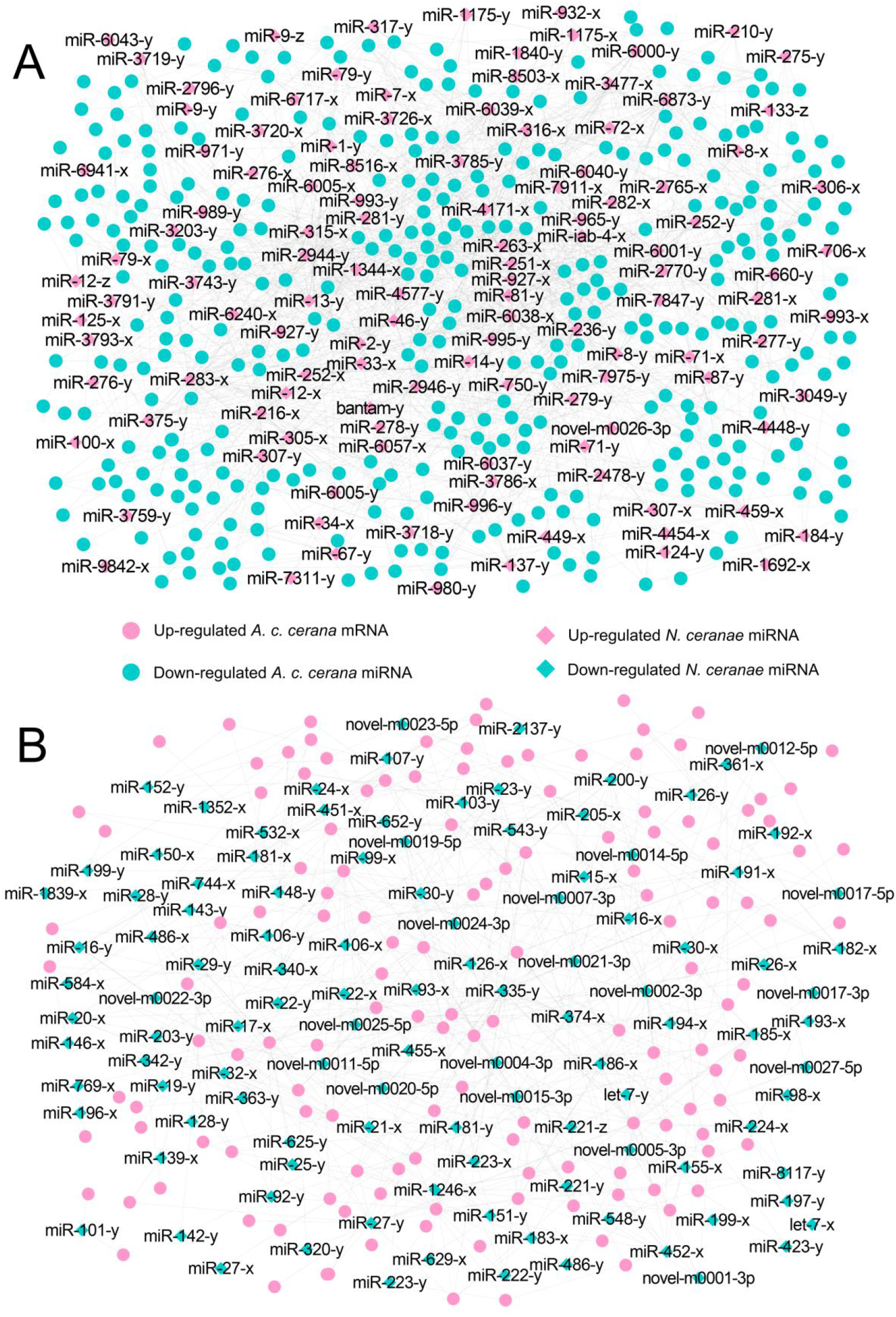
Regulatory networks of *N. ceranae* DEmiRNAs and target DEmRNAs in *A. c. cerana* workers’ midguts at 7 dpi. **(A)** Regulatory network between up-regulated miRNAs in microsporidian and down-regulated mRNAs in host. **(B)** Regulatory network down-regulated miRNAs in microsporidian and up-regulated mRNAs in host.

In AcCKM2 vs AcTM2 comparison group, 247 down- and 110 up-regulated mRNAs were putatively targeted by 110 up-regulated and 104 down-regulated miRNAs in NcCKMI vs NcTMI2 (**Figures 6, see also Table S2**). The above-mentioned 104 down-regulated mRNAs were engaged in 23 functional terms such as cellular process, membrane part, and catalytic activity; and 142 pathways including ubiquitin-mediated proteolysis, platinum drug resistance, and apoptosis-fly. Additionally, 110 up-regulated mRNAs were involved in 23 functional terms including metabolic process, binding, and catalytic activity; and 89 pathways such as quorum sensing, folate biosynthesis, and platinum drug resistance.

**FIGURE 6.**
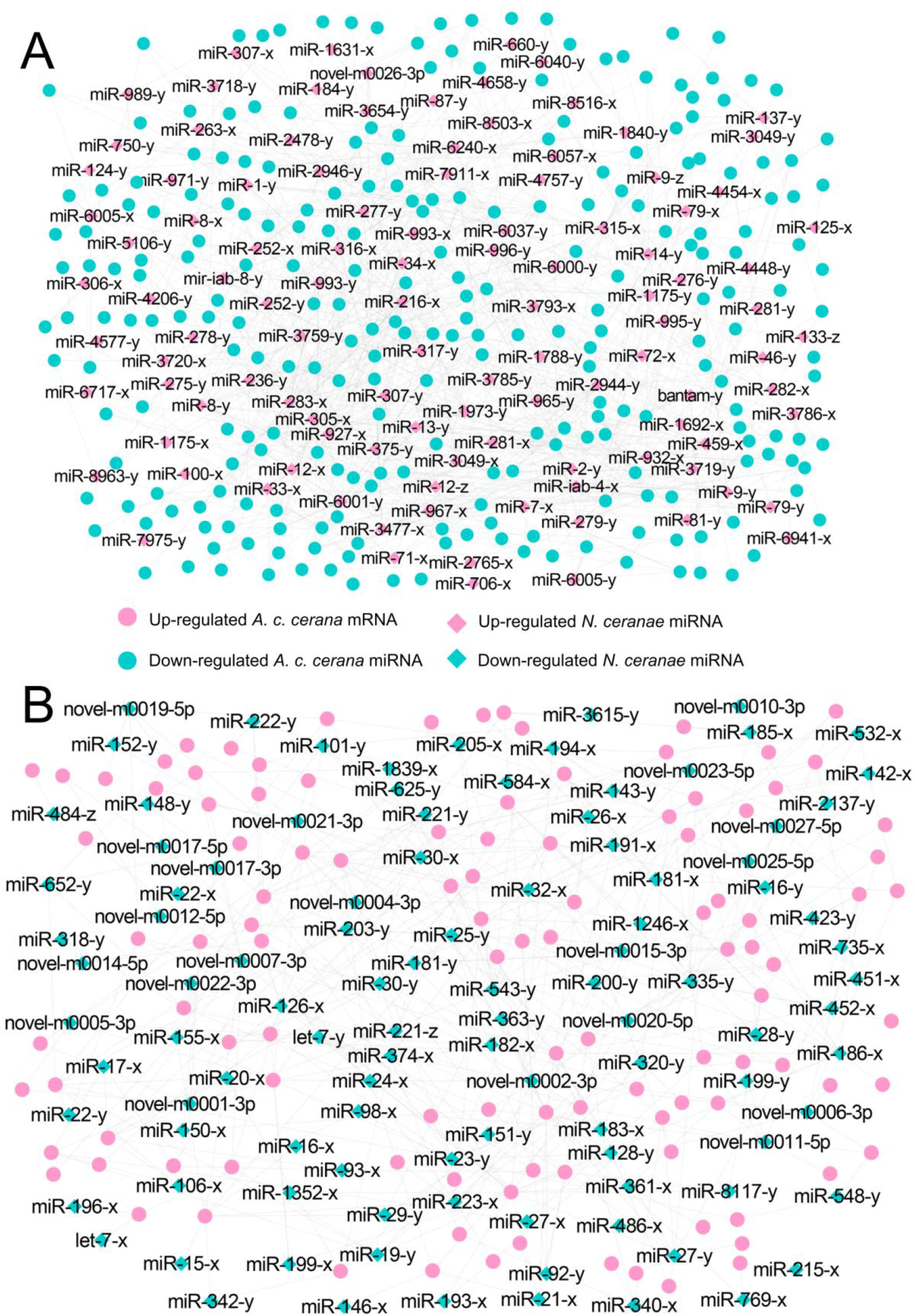
Regulatory networks of *N. ceranae* DEmiRNAs and target DEmRNAs in *A. c. cerana* workers’ midguts at 10 dpi. **(A)** Regulatory network between up-regulated miRNAs in microsporidian and down-regulated mRNAs in host. **(B)** Regulatory network down-regulated miRNAs in microsporidian and up-regulated mRNAs in host.

Further investigation showed that 62 up- and 42 down-regulated miRNAs shared by NcCKMI vs NcTMI1 and NcCKMI vs NcTMI2 comparison groups could potentially target 40 down-regulated and 15 common up-regulated mRNAs shared by AcCKM1 vs AcTM1 and AcCKM2 vs AcTM2 comparison groups (**Figures 7**).

**FIGURE 7.**
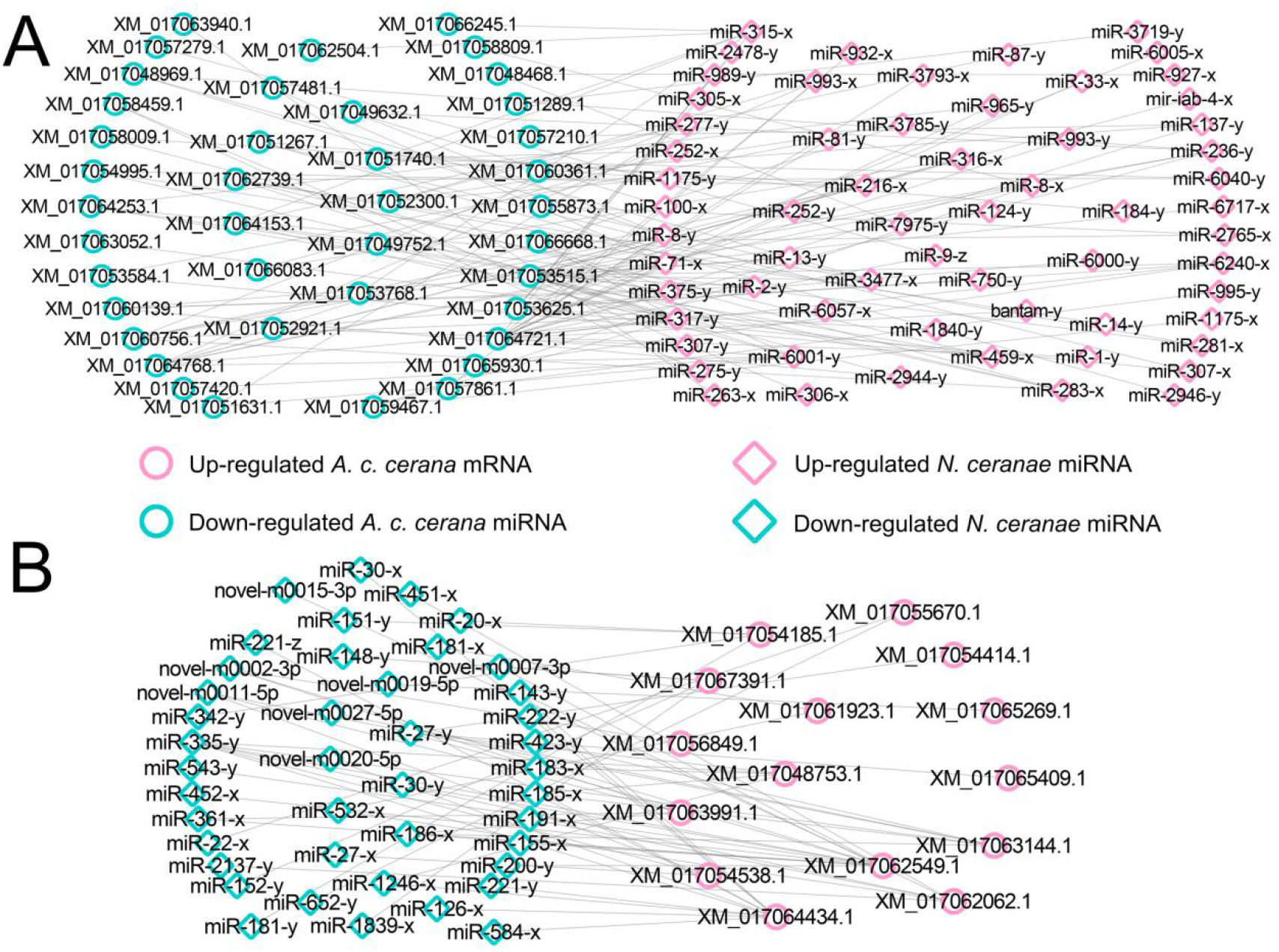
Regulatory networks of DEmiRNAs shared by NcCKMI vs NcTMI1 and NcCKMI vs NcTMI2 comparison groups and their targeted DEmRNAs shared by AcCKM1 vs AcTM1 and AcCKM2 vs AcTM2 comparison groups. **(A)** Network of common up-regulated miRNAs in microsporidian and common down-regulated mRNAs in host. **(B)** Network of shared down-regulated miRNAs in microsporidian and common up-regulated mRNAs in host.

**FIGURE 8.**
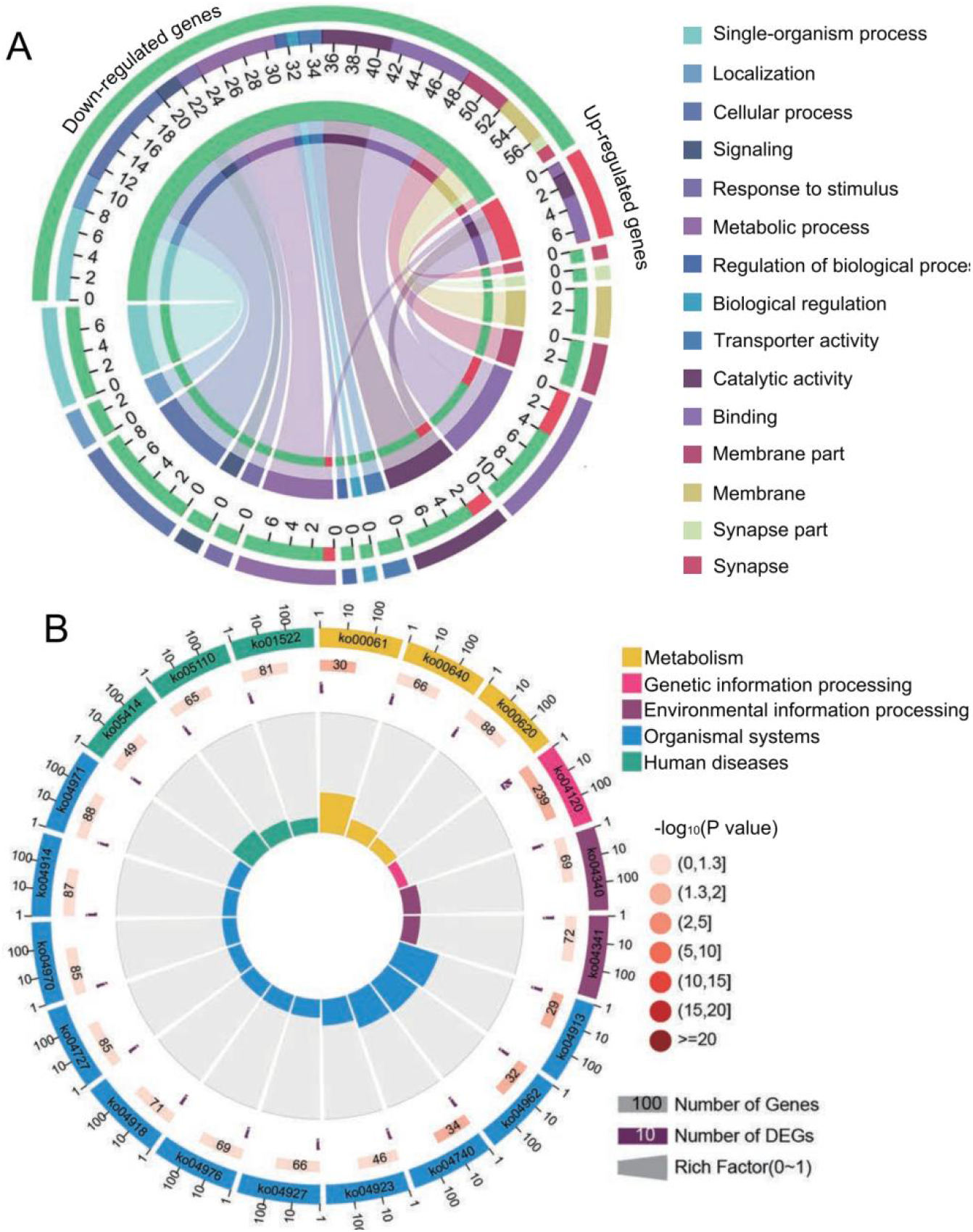
Database annotation of shared *A. c. cerana* DEmRNAs targeted by *N. ceranae* DEmiRNAs. **(A)** GO database annotation of DEmRNAs shared by AcCKM1 vs AcTM1 and AcCKM2 vs AcTM2. **(B)** KEGG database annotation of down-regulated mRNAs shared by AcCKM1 vs AcTM1 and AcCKM2 vs AcTM2.

Forty common down-regulated mRNAs in AcCKM1 vs AcTM1 and AcCKM2 vs AcTM2 targeted by 15 common up-regulated miRNAs in NcCKMI vs NcTMI1 and NcCKMI vs NcTMI2 were annotated to 18 GO terms; 12 mRNAs were engaged in biological process-associated categories, and the top three subcategories were cellular process, single-organism process, and metabolic process; four mRNAs were involved in cellular component-associated category, such as membrane part, membrane, cell, and cell part; ten mRNAs were engaged in molecular function-associated categories, among which the largest group was binding followed by catalytic activity and transporter activity. Additionally, the mRNAs mentioned above could be annotated to 70 pathways such as fatty acid biosynthesis, ovarian steroidogenesis, and aflatoxin biosynthesis.

### Investigation of *N. ceranae* DEmiRNAs and their target DEmRNAs associated with *A. c. cerana* immune and energy metabolism pathways

Further investigation indicated that 31 up-regulated miRNAs in NcCKMI vs NcTMI1 potentially targeted 12 down-regulated mRNAs in AcCKM1 vs AcTM1 (**Table S5**), which were involved in five immune-related pathways such as endocytosis, lysosomes, phagasome, Jak-STAT signaling pathway, and regulation of autophagy (**Figure 9A**). Comparatively, nine up-regulated miRNAs in NcCKMI vs NcTMI2 (**Table S5**) putatively targeted five down-regulated mRNAs in AcCKM2 vs AcTM2, which were engaged in three immune-related pathways including endocytosis, lysosomes, and regulation of autophagy (**Figure 9B**). In addition, the down-regulated miR-21-x (log2FC=-12.51, *P*<0.05) in NcCKMI vs NcTMI1 potentially targeted an up-regulated mRNA encoding NADH dehydrogenase [ubiquinone] 1 alpha subcomplex subunit 5 (XM_017057571.1, log2FC=1.28, *P*<0.01) in AcCKM1 vs AcTM1, which was relevant to oxidative phosphorylation, a key energy metabolism pathway.

**FIGURE 9.**
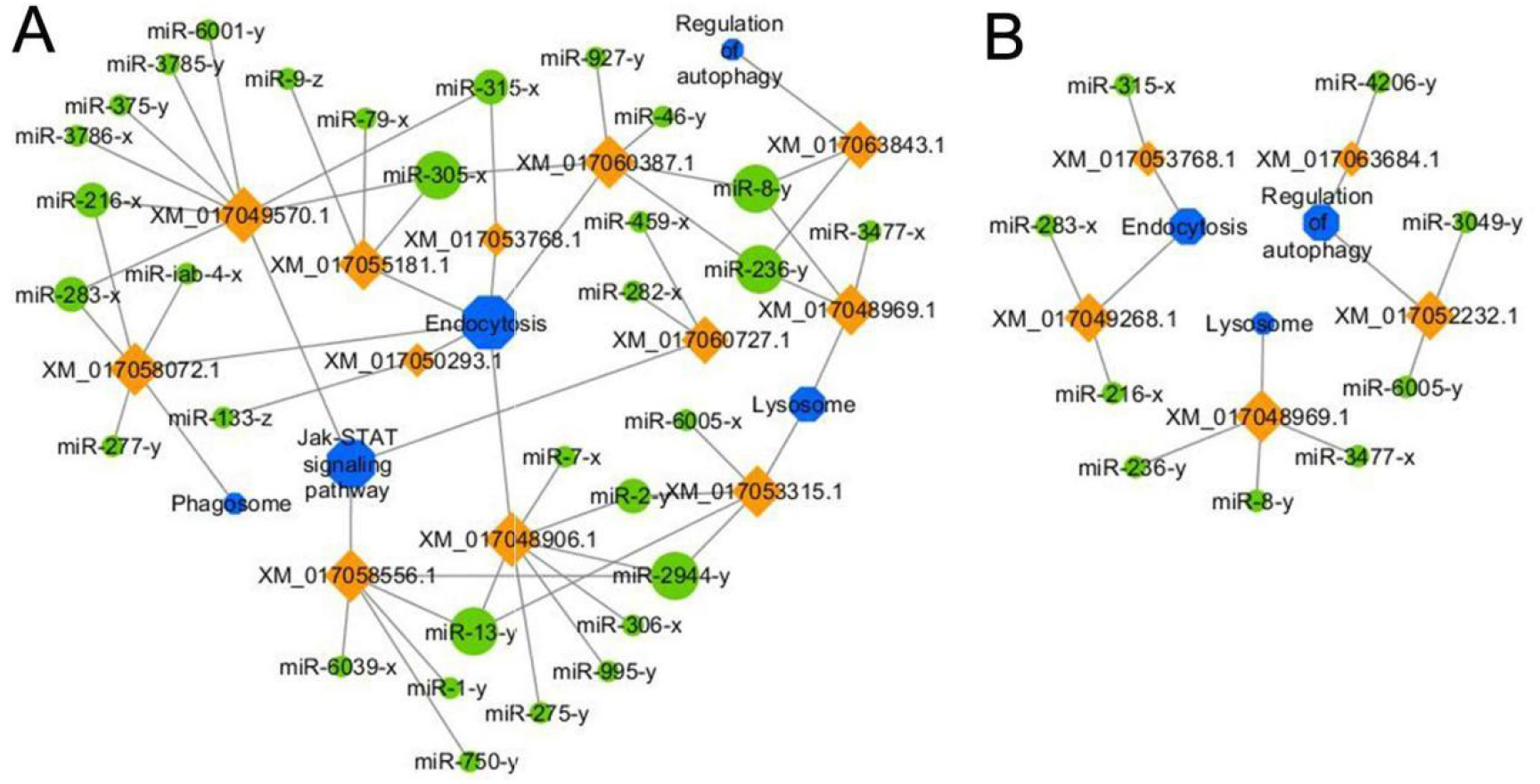
Regulatory networks of *N. ceranae* DEmiRNAs and their target DEmRNAs associated with *A. c. cerana* immune-related pathways. **(A)** Network of up-regulated miRNAs in NcCKMI vs NcTMI1 and immune-associated down-regulated mRNAs in AcCKM1 vs AcTM1. **(B)** Network of up-regulated miRNAs in NcCKMI vs NcTMI2 and immune-associated down-regulated mRNAs in AcCKM2 vs AcTM2.

### Validation of targeting relationship between *A. c. cerana* miRNAs and *N. ceranae* mRNAs and between *N. ceranae* miRNAs and *A. c. cerana* mRNAs

Following target prediction result in this study, two *A. c. cerana* DEmiRNAs (miR-8462-x and miR-676-y) and corresponding target DEmRNAs in *N. ceranae* (XM_002995668.1 and XM_002995068.1) as well as two *N. ceranae* DEmiRNAs (miR-2765-x and miR-6001-y) and corresponding target DEmRNAs in *A. c. cerana* (XM_017055873.1 and XM_017064721.1) were randomly selected for RT-qPCR validation, the result indicated that expression trends of above-mentioned four miRNAs and four mRNAs were in accordance with those in transcriptome data, and there was negative relationship between *A. c. cerana* DEmiRNAs and *N. ceranae* DEmRNAs as well as *N. ceranae* DEmiRNAs and *A. c. cerana* DEmRNAs (**Figure 10**), which confirmed the reliability of our transcriptome data and the targeting relationship between host and microsporidian.

**FIGURE 10.**
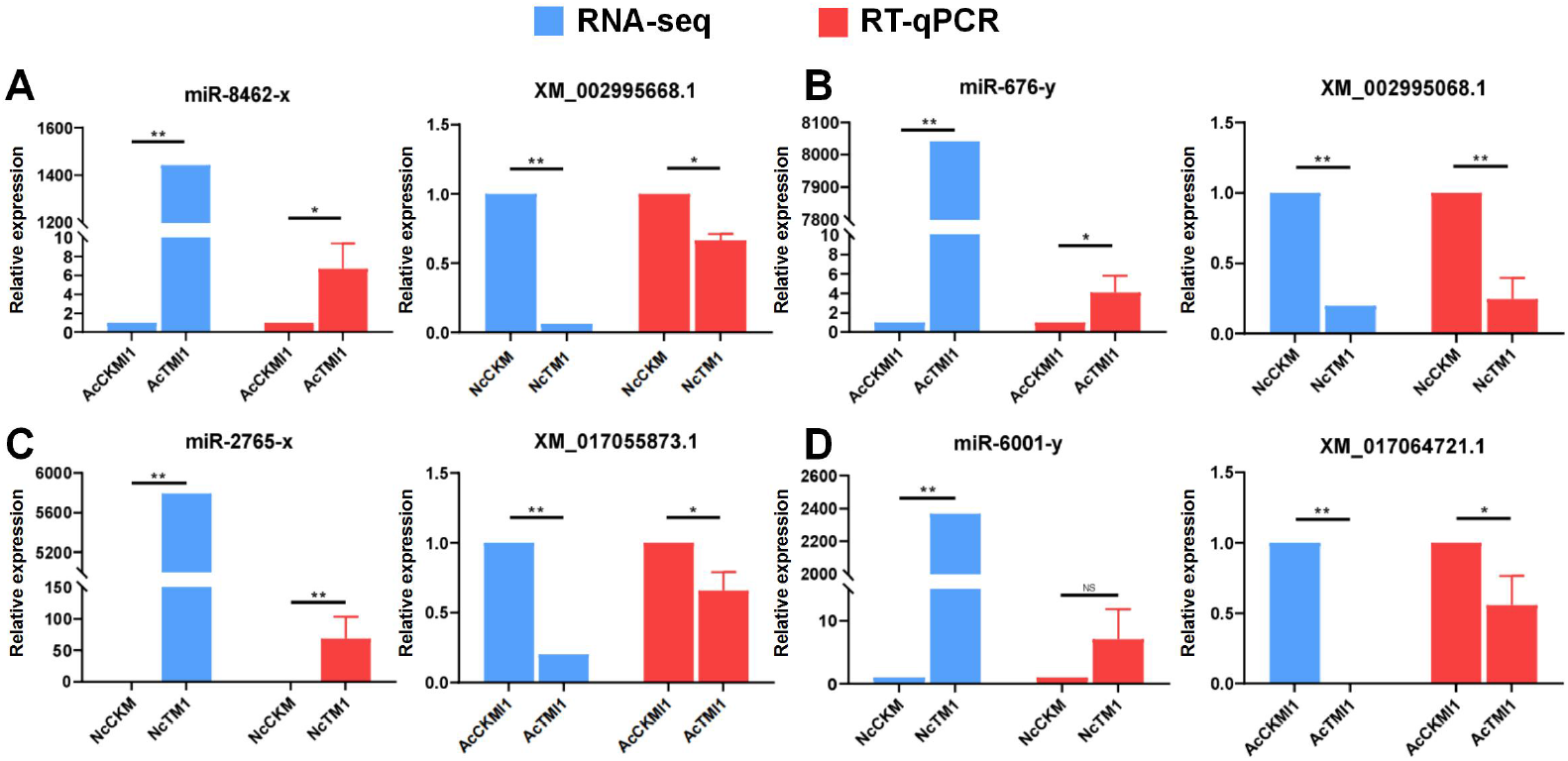
RT-qPCR validation of potential host miRNAs-microsporidian mRNAs and microsporidian miRNAs-host mRNAs targeting relationship. (A) RNA-seq and RT-qPCR results of *A. c. cerana* miR-8462-y and *N. ceranae* XM_002995668.1. (B) RNA-seq and RT-qPCR results of *A. c. cerana* miR-676-y and *N. ceranae* XM_002995068.1. (C) RNA-seq and RT-qPCR results of *N. ceranae* miR-2756-x and *A. c. cerana* XM_017055873.1. (D) RNA-seq and RT-qPCR results of *N. ceranae* miR-6001-y and *A. c. cerana* XM_017064721.1

## Discussion

Currently, cross-kingdom regulation between insects and fungal pathogens is poorly understood. As key regulators in regulation of gene expression and biological processes, miRNAs had been verified to mediate cross-kingdom regulation among animals, plants, and microorganisms (Rabuma et al., 2022; Halder et al., 2022). During the infection of *Solanum lycopersicum* and *Aedes albopictus, Botrytis cinerea* and *Beauveria bassiana* can synthesize and secrete sRNAs into the host cell via vesicles and hijacked the RNA interference mechanism by binding to the Argonaute 1 protein, which further silenced the expression of host immune genes (Weiberg et al., 2013; Cui et al., 2019). In cotton infected by *Verticillium dahliae*, the host-derived miR159 and miR166 were transported to the fungal mycelium and specifically silenced the fungal virulence factors isotrichomycin C-15 hydroxylase and Ca^2+^-dependent cysteine protease encoding genes, rendering cotton somewhat resistance to *V. dahliae* (Zhang et al., 2016). In the present study, we for the first time conducted comprehensive investigation of miRNA-mediated cross-kingdom regulation between *A. c. cerana* workers and *N. ceranae* utilizing previously obtained high-quality transcriptome data. It’s found that 97 down- and 60 up-regulated mRNAs in NcCKM vs NcTM1 were targeted by 8 up-and 6 down-regulated miRNAs in AcCKMI1 vs AcTMI1 (**Figure 1**), while 44 down- and 15 up-regulated mRNAs in NcCKM vs NcTM2 were targeted by seven up- and two down-regulated miRNAs in AcCKMI2 vs AcTMI2 (**Figure 2**); 343 down- and 138 up-regulated mRNAs in AcCKM1 vs AcTM1 were targeted by 121 up- and 112 down-regulated miRNAs in NcCKMI vs NcTMI1 (**Figure 5**), whereas 247 down- and 110 up-regulated mRNAs in AcCKM2 vs AcTM2 were targeted by 110 up- and 104 down-regulated miRNAs in NcCKMI vs NcTMI2 (**Figure 6**). The results was suggestive of potential targeting relationship between *A. c. cerana* DEmiRNAs and *N. ceranae* DEmRNAs as well as *N. ceranae*DEmiRNAs and *A. c. cerana* DEmRNAs.

### *A. c. cerana* DEmiRNAs putatively regulate glycolysis/glyconeogenesis and virulence factors in *N. ceranae*

With an extremely reduced genome, *N. ceranae* is unable to produce ATP via the tricarboxylic acid (TCA) cycle and oxidative phosphorylation, thus highly depending on host cell-produced energy; however, *N. ceranae* retains the intact glycolysis pathway, which is active primarily in the spore stage (Dolgikh et al., 1997, He et al., 2020; Hoch et al., 2002). In this current work, host miR-676-y (log_2_FC=12.97, *P*<0.01) was detected to target XM_002996090.1 (log_2_FC=-6.89, *P*<0.01) associated with the glycolysis/gluconeogenesis pathway in *N. ceranae* at 7 dpi (**Figure 5**); additionally, the host miR-60-y (log_2_FC=14.32, *P*<0.01) was observed to target XM_002995904.1 (log_2_FC=3.78, *P*<0.01) in *N. ceranae* at 10 dpi (**Figure 5**). The results implied that miR-676-y and miR-60-y was potentially employed by host to down-regulate the expression of associated genes and impact glycolysis/gluconeogenesis pathway, which may further inhibit the energy metabolism of *N. ceranae*.

Microsporidian cells are simplified and lack mitochondria, instead contain a genome-less organelle named mitosome, which appears to function in iron-sulfur biochemistry but not oxidative phosphorylation (Burri et al., 2006). The thick spore wall of microsporidian is composed of proteins and chitins, which helps the microsporidia survive in various harsh environments (Vávra, 1976). Spore wall proteins (SWPs) have also been verified to interact with host cells and participate in the infection process of microsporidia (Jaroenlak et al., 2018; Yang et al., 2018). NbSWP5 was reported to be involved in the germination of *N. bombycis* spores by interaction with the polar tube proteins and protect spores from phagocytic uptake by cultured insect cells (Cai et al., 2011; Li et al., 2012). The oral ingestion of dsRNAs corresponding to SWP12 caused significant reduction of the *N. ceranae* spore load in *A. m. ligustica* workers’ midguts, and at the meanwhile improved host immune defense and extent the host lifespan (He et al., 2021). In *C. elegans*, miR-60 was exclusively expressed in the intestinal tissue, and seemed to directly modulate the endocytosis machinery and zip-10-mediated innate immune systems and coordinate the expression of genes engaged in the cellular homeostasis, further promoting adaptive response against the long-term mild oxidative stress (Kato et al., 2016). Here, miR-60-y was found to significantly up-regulate (log_2_FC=10.87, *P*<0.01) in *A. c. cerana* worker’s midgut and target the mRNA (XM_002996592.1, log_2_FC=-2.24, *P*<0.01) encoding SWP12 in *N. ceranae* at 7 dpi, indicating that miR-60-y may be utilized by host to suppress the *N. ceranae* infection via negative regulation of the expression of *SWP12*. Together, these results demonstrated that the aforementioned three DEmiRNAs in *A. cerana* workers were potentially activated to target and inhibit genes associated with glycolysis/glyconeogenesis and SWPs during the *N. ceranae* infection.

In view of that effective overexpression of fungal miRNA can be achieved using corresponding mimics. For example, Chakrabarti et al. (2020) found that chemically synthesized mimics of erythrocytic miR-150-3p and miR-197-5p were loaded into erythrocytes and subsequently used for invasion by the parasite, growth of the *Plasmodium falciparum* was hindered in miRNA-loaded erythrocytes, and both micronemal secretion and *Apicortin* expression were reduced in miRNA-loaded erythrocytes. When treating RAW 264.7 cells infected by *Leishmania major* promastigotes (MRHO/IR/75/ER) with miR-15a mimic and (or) miR-155 inhibitor, Gholamrezaei et al. (2020) observed that apoptosis of macrophages was increased and at the meantime parasite burden was reduced. At present, cultured cells for *N. ceranae* is still lacking, limiting in vitro functional study of *N. ceranae* genes. However, *N. ceranae*-infected honey bee model had been constructed and applied to functional study on genes associated with fungal proliferation. Huang et al. (2019) previously injected siRNA-*Dicer* into *A. mellifera* workers infected by *N. ceranae*, observed the down-regulation of genes encoding two virulence factors including ABC transporter and hexokinase, and found that the spore load was significantly reduced, indicating that silencing of *Dicer* gene could suppress the proliferation of *N. ceranae*.Recently, Kim et al. (2020) conducted dsRNA-based RNAi of mitosome-related genes in *N. ceranae*, and discovered that the microsporidian proliferation was inhibited, while the host survival rate was increased. To clarify function of above-mentioned genes targeted by host miR-676-y and miR-60-y, siRNA- or dsRNA-based RNAi of these targets will be conducted in the near future.

### *N. ceranae* DEmiRNAs potentially regulate energy mechanism and immune response of *A. c. cerana* workers

Microsporidia are a group of obligate intracellular parasites that lack mitochondria, thus microsporidia evolve unique strategies to acquire nutrients from host cells and manipulate host metabolism (Yang et al., 2018). By analyzing the miRNA response of *A. mellifera* workers to *N. ceranae* infection, Evans and Huang (2018) revealed that *N. ceranae* DEmiRNAs may regulate expression of host genes relevant to purine metabolism, pyrimidine metabolism, and oxidative phosphorylation. In this study, the miR-21-x (log_2_FC=-12.51, *P*<0.05) was significantly down-regulated and potentially target the up-regulated mRNA (XM_017057571.1, log_2_FC=1.13, *P*<0.01) involved in host oxidative phosphorylation at 7 dpi (**Figure 9; Table S5**), indicative of the activation of host energy mechanism microsporidian. It’s inferred that oxidative phosphorylation in host cells was induced to activation by *N. ceranae* to provide more ATP for microsporidian proliferation.

Similar to other insects, honey bees have evolved an innate immune instead of adaptive immune to fight against foreign invaders (Evans et al., 2006). Insect innate immune is consisted of three main processes: (1) pathogen recognition; (2) signal transduction; (3) regulation of immune signaling pathways, expression of downstream genes, and final immune responses (Naitza and Ligoxygakis, 2004). Insect immune system could be divided into cellular and humoral immune (McMenamin et al., 2018), however, these two kinds of immune cannot be strictly separated since both are interrelated and act together to generate highly effective defense (Li et al., 2018). The humoral immunity senses a wide range of pathogens through distinct pattern recognition receptors (PRRs), and the engagement of these PRRs induces the activation of specific signaling pathways, leading to the expression of antimicrobial genes (McMenamin et al., 2018). JAK/STAT signaling pathway plays a central role in immune response of bee individual (McMenamin et al., 2018). In silkworms, weak response of JAK/STAT signaling pathway could be induced by the fungal infection (Liu et al., 2015). Antúnez et al. (2009) detected that the immune response of *A. m. ligustica* workers was suppressed at 7 dpi with *N. ceranae*, and the expression of genes encoding antimicrobial peptides and immune-related enzymes in *N. ceranae*-infected workers was significantly down-regulated as compared with un-infected workers. Here, a total of 15 up-regulated miRNAs in NcCKMI vs NcTMI1 comparison group were found to target three down-regulated mRNAs enriched in Jak/STAT signaling pathway in AcCKM1 vs AcTM1 comparison group. In addition, miR-216-x (NcCK vs NcTMI1: log_2_FC=14.92, *P*<0.01; NcCK vs NcTMI2: log_2_FC=15.10, *P*<0.01), miR-236-y (NcCK vs NcTMI1: log_2_FC=12.98, *P*<0.01; NcCK vs NcTMI2: log_2_FC=12.96, *P*<0.01), miR-283-x (NcCK vs NcTMI1: log_2_FC=9.41, *P*<0.05; NcCK vs NcTMI2: log_2_FC=9.08, *P*<0.05), miR-315-x (NcCK vs NcTMI1: log_2_FC=17.11, *P*<0.01; NcCK vs NcTMI2: log_2_FC=16.33, *P*<0.01), miR-3477-x (NcCK vs NcTMI1: log_2_FC=22.22,*P*<0.01; NcCK vs NcTMI2: log_2_FC=21.73, *P*<0.01), and miR-8-y (NcCK vs NcTMI1: log_2_FC=9.68, *P*<0.05; NcCK vs NcTMI2: log_2_FC=9.54, *P*<0.05) were up-regulated in both NcCKM vs NcTM1 and NcCKM vs NcTM2 comparison groups and potentially targeted corresponding down-regulated mRNA relevant to immune pathways such as XM_017052232.1 (log_2_FC=-11.13, *P*<0.01) and XM_017049268.1 (log_2_FC=-12.02, *P*<0.01) (**Figure 9; Table S5**).

The *A. mellifera* genome contains genes encoding most of members in insect immune pathways, including orthologues of genes engaged in the autophagy and endocytosis (McMenamin et al., 2018). Endocytosis is a dynamic process that regulates various signaling pathways in both positive and negative manners(Papagiannouli, 2022). Lysosome has antifungal and antiviral activities and dissolves pathogenic proteins delivered by endocytosis and phagocytosis (Samaranayake et al., 2001). Here, miR-315-x (NcCK vs NcTMI1: log_2_FC=17.11, *P*<0.01; NcCK vs NcTMI2: log_2_FC=16.33, *P*<0.01) shared by NcCK vs NcTMI1 and NcCK vs NcTMI2 comparison groups, were observed to down-regulate and target the mRNAs XM_017053768.1 (NcCK vs NcTMI1: log_2_FC=-9.97, *P*<0.01; NcCK vs NcTMI2: log_2_FC=-9.44, *P*<0.01) which was relative to endocytosis; additionally, miR-3477-x (NcCK vs NcTMI1: log_2_FC=22.22, *P*<0.01; NcCK vs NcTMI2: log_2_FC=21.73, *P*<0.01), miR-8-y (NcCK vs NcTMI1: log_2_FC=9.68, *P*<0.05; NcCK vs NcTMI2: log_2_FC=9.54, *P*<0.05), and miR-236-y (NcCK vs NcTMI1: log_2_FC=12.98, *P*<0.01; NcCK vs NcTMI2: log_2_FC=12.96, *P*<0.01) were up-regulated in both NcCK vs NcTMI1 and NcCK vs NcTMI2 comparison groups and putatively targeted the same one mRNA XM_017048969.1 (NcCK vs NcTMI1: log_2_FC=-11.47, *P*<0.01; NcCK vs NcTMI2: log_2_FC=-10.21, *P*<0.05), which was relative to lysosome. These findings suggested that *N. ceranae* was likely to up-regulate the expression of above-mentioned DEmiRNAs to weaken the host humoral and cellular immune during the infection.

Taken together, we for the first time deciphered the landscape of DEmRNA-mediated cross-kingdom regulation between *A. c. cerana* workers and *N. ceranae*, and unraveled that host DEmiRNAs probably regulate glycolysis/glyconeogenesis and virulence factors during the *N. ceranae* infection to suppress the microsporidian proliferation, whereas *N. ceranae* DEmiRNAs were likely to enhance host energy mechanism and inhibit host immune responses to facilitate microsporidian invasion. Findings in this current work lay a foundation for exploring the molecular mechanism underlying cross-kingdom regulation between *A. c. cerana* workers and *N. ceranae*, provide valuable insights into Asian honey bee-microsporidian interaction, and offer potential targets for bee nosemosis control.

## Acknowledgements

This work was financially supported by the National Natural Science Foundation of China

(32172792), the Earmarked fund for CARS-44-KXJ7 (CARS-44-KXJ7), the Master Supervisor Team Fund of Fujian Agriculture and Forestry University (Rui Guo), the Scientific Research Project of College of Animal Sciences (College of Bee Science) of Fujian Agriculture and Forestry University (Rui Guo), the National Undergraduate Innovation and Entrepreneurship Training Program (Kuihao Zhang), and the Fund for Excellent Master Dissertation of Fujian Agriculture and Forestry University (Xiaoxue Fan).

## REFERENCES

Allen, E., Xie, Z., Gustafson, A. M., and Carrington, J. C. (2005). microRNA-directed phasing during trans-acting siRNA biogenesis in plants. Cell. 121(2), 207–221. doi: 10.1016/j.cell.2005.04.004

Antúnez, K., Martín-Hernández, R., Prieto, L., Meana, A., Zunino, P., and Higes, M. (2009). Immune suppression in the honey bee *(Apis mellifera)* following infection by *Nosema ceranae* (Microsporidia). Environ. Microbiol. 11(9), 2284–2290. doi: 10.1111/j.1462-2920.2009.01953.x

Arias, M. C., and Sheppard, W. S. (2005). Phylogenetic relationships of honey bees (Hymenoptera:Apinae: Apini) inferred from nuclear and mitochondrial DNA sequence data. Mol. Phylogene. Evol. 37(1), 25–35. doi: 10.1016/j.ympev.2005.02.017

Bartel D. P. (2004). MicroRNAs: genomics, biogenesis, mechanism, and function. Cell. 116(2), 281–297. doi: 10.1016/s0092-8674(04)00045-5

Burri, L., Williams, B. A., Bursac, D., Lithgow, T., and Keeling, P. J. (2006). Microsporidian mitosomes retain elements of the general mitochondrial targeting system. Proc. Natl. Acad. Sci. USA., 103(43), 15916–15920. doi: https://doi.org/10.1073/pnas.0604109103

Chaimanee, V., Chantawannakul, P., Chen, Y., Evans, J. D., and Pettis, J. S. (2012). Differential expression of immune genes of adult honey bee (*Apis mellifera*) after inoculated by *Nosema ceranae*. J. Insect Physiol. 58(8), 1090–1095. doi: 10.1016/j.jinsphys.2012.04.016

Chakrabarti, M., Garg, S., Rajagopal, A., Pati, S., and Singh, S. (2020). Targeted repression of Plasmodium apicortin by host microRNA impairs malaria parasite growth and invasion. Disease models & mechanisms, 13(6), dmm042820. doi: https://doi.org/10.1242/dmm.042820

Chen, C., Eldein, S., Zhou, X., Sun, Y., Gao, J., Sun, Y., Liu, C., and Wang, L. (2018). Immune function of a Rab-related protein by modulating the JAK-STAT signaling pathway in the silkworm, *Bombyx mori*. Arch. Insect Biochem. Physiol. 97(1). doi: 10.1002/arch.21434

Chen, D., Chen, H., Du, Y., Zhou, D., Geng, S., Wang, H., et al. (2019). Genome-Wide Identification of Long Non-Coding RNAs and Their Regulatory Networks Involved in *Apis mellifera ligustica* Response to *Nosema ceranae* Infection. Insects, 10(8), 245. doi: https://doi.org/10.3390/insects10080245

Chen, D. F., Guo, R., Xiong, C. L., Liang, Q., Zheng, Y. Z., Xu, X. J. et al. (2017). Transcriptomic analysis of Ascosphaera apis stressing larval gut of *Apis mellifera ligustica* (Hyemenoptera: Apidae), Acta Entomologica Sinica, 60(4), 401–411. (in Chinese)

Chen, D., Du, Y., Chen, H., Fan, Y., Fan, X., Zhu, Z., et al. (2019). Comparative identification of micrornas in *Apis cerana cerana* workers’ midguts in responseto *Nosema ceranae* invasion. Insects. 10(9), 258. doi: 10.3390/insectmi10090258

Cornman, R. S., Chen, Y. P., Schatz, M. C., Street, C., Zhao, Y., Desany, B., et al. (2009). Genomic analyses of the microsporidian *Nosema ceranae*, an emergent pathogen of honey bees. PLoS Pathog. 5(6), e1000466. doi: 10.1371/journal.ppat.1000466

Cui, C., Wang, Y., Liu, J., Zhao, J., Sun, P., and Wang, S. (2019). A fungal pathogen deploys a small silencing RNA that attenuates mosquito immunity and facilitates infection. Nat. Commun. 10(1), 4298. doi: 10.1038/s41467-019-12323-1

Cuomo, C. A., Desjardins, C. A., Bakowski, M. A., Goldberg, J., Ma, A. T., Becnel, J. J., et al. (2012). Microsporidian genome analysis reveals evolutionary strategies for obligate intracellular growth. Genome Res. 22(12), 2478–2488. doi: 10.1101/gr.142802.112

Dermauw, W., and Van Leeuwen, T. (2014). The ABC gene family in arthropods: comparative genomics and role in insecticide transport and resistance. Insect Biochem. Mol. Biol. 45, 89–110. doi: 10.1016/j.ibmb.2013.11.001

Diao, Q. Y., Sun, L. X., Zheng, H. J., Zeng, Z. J., Wang, S. Y., Xu, S. F., et al. (2018). Genomic and transcriptomic analysis of the Asian honey bee Apis cerana provides novel insights into honey bee biology. Sci. Rep. 8(1): 822. doi: 10.1038/s41598-017-17338-6

Dolgikh, V. V., Sokolova, J. J., and Issi, I. V. (1997). Activities of enzymes of carbohydrate and energy metabolism of the spores of the microsporidian, *Nosema grylli*. J. Eukaryot. Microbiol. 44(3), 246–249. doi:10.1111/j.1550-7408.1997.tb05707.x

Du, Y., Fan, X. X., Jiang, H. B., Wang, J., Feng, R. R., Zhang, W. D., et al. (2021). MicroRNA-mediated cross-kingdom regulation of *Apis mellifera ligustica* worker to *Nosema ceranae*. Scientia Agricultura Sinica. 54(8), 1805–1820. (in Chinese)

Eiri, D.M., Suwannapong, G., Endler, M., Nieh, J.C. (2015) *Nosema ceranae* Can Infect Honey Bee Larvae and Reduces Subsequent Adult Longevity. PLoS One. 10(5), e0126330. doi: 10.1371/journal.pone.0126330

Evans, J. D., Aronstein, K., Chen, Y. P., Hetru, C., Imler, J. L., Jiang, H., et al. (2006). Immune pathways and defence mechanisms in honey bees *Apis mellifera*. Insect Mol. Biol. 15(5), 645–656. doi: 10.1111/j.1365-2583.2006.00682.x

Evans, J. D., and Huang, Q. (2018). Interactions Among Host-Parasite MicroRNAs During *Nosema ceranae* Proliferation in *Apis mellifera*. Frontiers in microbiology, 9, 698. doi: https://doi.org/10.3389/fmicb.2018.00698

Fan, X. X., Du, Y., Zhang, W. D., Wang, J., Jiang, H. B., Fan, Y. C. et al. (2021). Omics analysis of *Nosema ceranae* miRNAs involved in gene expression regulation in the midguts of *Apis mellifera ligustica* workers and their regulatory networks. Acta Entomologica Sinica, 64(2), 187–204. (in Chinese)

Fan, Y., Wang, J., Yu, K., Zhang, W., Cai, Z., Sun, M., et al. (2022). Comparative Transcriptome Investigation of *Nosema ceranae* Infecting Eastern Honey Bee Workers. Insects. 13(3), 241.

Ferguson, S., and Lucocq, J. (2019). The invasive cell coat at the microsporidian Trachipleistophora hominis-host cell interface contains secreted hexokinases. MicrobiologyOpen. 8(4), e00696. doi: 10.1002/mbo3.696

Franzen, C. (2005). How do microsporidia invade cells?. Folia Parasitol (Praha). 52(1-2), 36–40. doi: 10.14411/fp.2005.005

Fries, I., Feng, F., Silva, A. D., Slemenda, S. B., and Pieniazek, N. J. (1996). *Nosema ceranae* n. sp. (Microspora, Nosematidae), morphological and molecular characterization of a microsporidian parasite of the Asian honey bee *Apis cerana* (Hymenoptera, Apidae). Europ. J. Protistology. 32, 356–365. doi: 10.1016/S0932-4739(96)80059-9

Fu, Z. M., Zhou, D. D., Chen, H. Z., Geng, S. H., Chen D. F., Zheng, Y. Z. et al. (2020). Analysis of highly expressed genes in *Apis cerana cerana* workers’ midguts responding to *Nocema ceranae* stress. Journal of Sichuan University(Natural Science Edition), 57(01), 191–198. (in Chinese)

Fu, Z. M., Zhou, D. D., Chen, H. Z., Geng, S. H., Chen, D. F., Zheng, Y. Z., et al. (2020). Analysis of highly expressed genes in *Apis cerana cerana* workers’ midguts responding to *Nosema ceranae* stress. J. Sichuan Univ. (Nat. Sci. Ed.), 57, 191–198. (in Chinese)

Fu, Z. M., Chen, H. Z., Liu, S.Y., Zhu, Z. W., Fan, X. X., Fan, Y. C., et al. (2019). Immune Responses of *Apis mellifera ligustia* to *Nosema ceranae* Stress. Scientia Agricultura Sinica, 52(17), 3069–3082. (in Chinese)

Gallai, N., Salles, J. M., Settele, J., Vaissière, B. E. (2009). Economic valuation of the vulnerability of world agriculture confronted with pollinator decline. Ecol. Econ. 68, 810–821. doi: 10.1016/j.ecolecon.2008.06.014

Garofalo, M., and Croce, C. M. (2011). microRNAs: Master regulators as potential therapeutics in cancer. Annu. Rev. Pharmacol. Toxicol., 51, 25–43. doi: https://doi.org/10.1146/annurev-pharmtox-010510-100517

Geng, S. H., Shi, C. Y., Fan, X. X., Wang, J., Zhu, Z. W., Jiang, H. B. et al. (2020a). The Mechanism Underlying MicroRNAs-Mediated *Nosema ceranae* Infection to *Apis mellifera ligustica* Worker, Scientia Agricultura Sinica, 53(15), 3187–3204. (in Chinese)

Geng, S. H., Shi, C. Y., Fan, X. X., Wang, J., Zhu, Z. W., Jiang, H. B., et al. (2020b). Revealing the mechanism underlying *Nosema ceranae* infection of *Apis mellifera ligustica* based on investigation of differential expression profile and regulatory network of microRNAs. Scientia Agricultura Sinica. (in Chinese)

Geng, S. H., Zhou, D. D., Fan, X. X., Jiang, H. B., Zhu, Z. W., Wang, J., et al. (2020c). Transcriptomic analysis reveals the molecular mechanism underlying *Nosema ceranae* infection of *Apis mellifera ligustica*. Scientia Agricultura Sinica, 63(03), 294–308. (in Chinese)

George, P. R. (2008). Encyclopedia of Genetics, Genomics, Proteomics and Informatics. Springer Netherlands. 752–753.

Gholamrezaei, M., Rouhani, S., Mohebali, M., Mohammadi-Yeganeh, S., Haji Molla Hoseini, M., Haghighi, A., et al. (2020). MicroRNAs expression induces apoptosis of macrophages in response to leishmania major (MRHO/IR/75/ER): An In-Vitro and In-Vivo Study. Iranian journal of parasitology, 15(4), 475–487. doi: https://doi.org/10.18502/ijpa.v15i4.4851

Guo, R., Chen, D. F., Xiong, C. L., Hou, C. S., Zheng, Y. Z., Fu, Z. M., et al. (2018). First identification of long non-coding RNAs in fungal parasite *Nosema ceranae*. Apidologie. 8(49), 660–670. doi: 10.1007/s13592-018-0593-z

Halder, L. D., Babych, S., Palme, D. I., Halder, L. D., Babych, S., Palme, D. I., et al. (2022). Candida albicans induces cross-kingdom mirna trafficking in human monocytes to promote fungal growth. mBio, 13(1), e0356321. Advance online publication. doi: https://doi.org/10.1128/mbio.03563-21

Hamel, L. P., Nicole, M. C., Duplessis, S., and Ellis, B. E. (2012). Mitogen-activated protein kinase signaling in plant-interacting fungi: distinct messages from conserved messengers. Plant. Cell. 24(4), 1327–1351. doi: 10.1105/tpc.112.096156.

He, N., Zhang, Y., Duan, X. L., Li, J. H., Huang, W. F., Evans, J. D., et al. (2021). RNA interference-mediated knockdown of genes encoding spore wall proteins confers protection against *Nosema ceranae* infection in the European honey bee, *Apis mellifera*. Microorganisms. 9(3), 505. doi: 10.3390/microorganisms9030505

He, Q., Luo, J., Xu, J. Z., Wang, C. X., Meng, X. Z., Pan, G. Q., et al. (2020). Morphology and transcriptome analysis of *Nosema bombycis* sporoplasm and insights into the initial infection of microsporidia. mSphere. 5(1), e00958–19. doi: 10.1128/mSphere.00958-19

Heinz, E., Hacker, C., Dean, P., Mifsud, J., Goldberg, A. V., Williams, T. A., et al. (2014). Plasma membrane-located purine nucleotide transport proteins are key components for host exploitation by microsporidian intracellular parasites. PLoS Pathog. 10(12), e1004547. doi: 10.1371/journal.ppat.1004547

Heinz, E., Williams, T. A., Nakjang, S., Noël, C. J., Swan, D. C., Goldberg, A. V., et al. (2012). The genome of the obligate intracellular parasite *Trachipleistophora hominis:* new insights into microsporidian genome dynamics and reductive evolution. PLoS Pathog. 8(10), e1002979. doi: 10.1371/journal.ppat.1002979

Hepburn, H. R., and Radloff, S. E. (2011). honey bees of Asia. Springer

Hoch, G., Schafellner, C., Henn, M. W., and Schopf, A. (2002). Alterations in carbohydrate and fatty acid levels of *Lymantria dispar* larvae caused by a microsporidian infection and potential adverse effects on a co-occurring endoparasitoid, *Glyptapanteles liparidis*. Arch. Insect. Biochem. Physiol. 50(3), 109–120. doi:10.1002/arch.10030

Huang, Q., and Evans, J. D. (2016). Identification of microRNA-like small RNAs from fungal parasite *Nosema ceranae*. J. Invertebr. Pathol. 133, 107–109. doi: 10.1016/j.jip.2015.12.005

Huang, Q., Chen, Y. P., Wang, R. W., Cheng, S., Evans, J. D. (2016b). Host-parasite interactions and purifying selection in a Microsporidian parasite of honey bees. PloS one, 11(2): e0147549. doi: 10.1371/journal.pone.0147549

Huang, Q., Li, W., Chen, Y., Retschnig-Tanner, G., Yanez, O., Neumann, P., et al. (2019). Dicer regulates *Nosema ceranae* proliferation in honey bees. Insect Mol. Biol. 28(1), 74–85.

Jaroenlak, P., Boakye, D. W., Vanichviriyakit, R., Williams, B., SritunyaluCKMIana, K., and Itsathitphaisarn, O. (2018). Identification, characterization and heparin binding capacity of a spore-wall, virulence protein from the shrimp microsporidian, *Enterocytozoon hepatopenaei* (EHP). Parasit vectors. 11(1), 177. doi:10.1186/s13071-018-2758-z

Kanehisa, M., and Goto, S. (2000). KEGG: kyoto encyclopedia of genes and genomes. Nucleic Acids Res. 28(1), 27–30. doi: 10.1093/nar/28.1.27

Katinka, M. D., Duprat, S., Cornillot, E., Méténier, G., Thomarat, F., Prensier, G., et al. (2001). Genome sequence and gene compaction of the eukaryote parasite *Encephalitozoon cuniculi*. Nature. 414(6862), 450–453. doi: 10.1038/35106579

Kato, M., Kashem, M. A., and Cheng, C. (2016). An intestinal microRNA modulates the homeostatic adaptation to chronic oxidative stress in C. elegans. Aging. 8(9), 1979–2005. doi: 10.18632/aging.101029

Kim, I. H., Kim, D. J., Gwak, W. S., and Woo, S. D. (2020). Increased survival of the honey bee *Apis mellifera* infected with the microsporidian *Nosema ceranae* by effective gene silencing. Arch. Insect Biochem. Physiol. 105(4), e21734. doi:10.1002/arch.21734

Leclerc, V., and Reichhart, J. M. (2004). The immune response of *Drosophila melanogaster*. Immunol. Rev. 198, 59–71. doi: 10.1111/j.0105-2896.2004.0130.x

Li, W., Chen, Y., and Cook, S. C. (2018). Chronic *Nosema ceranae* infection inflicts comprehensive and persistent immunosuppression and accelerated lipid loss in host *Apis mellifera* honey bees. Int. J. Parasitol. 48(6), 433–444. doi: 10.1016/j.ijpara.2017.11.004

Li, Z., Pan, G., Li, T., Huang, W., Chen, J., Geng, L., Yang, D., Wang, L., and Zhou, Z. (2012). SWP5, a spore wall protein, interacts with polar tube proteins in the parasitic microsporidian *Nosema bombycis*. Eukaryot. Cell. 11(2), 229–237. doi:10.1128/EC.05127-11

Liu, H., Li, M., Cai, S., He, X., Shao, Y., and Lu, X. (2016). Ricin-B-lectin enhances microsporidia *Nosema bombycis* infection in *BmN* cells from silkworm *Bombyx mori*. Acta biochim. Biophys. Sin. 48(11), 1050–1057. doi: 10.1093/abbs/gmw093

Liu, H., Wang, X., Wang, H. D., Wu, J., Ren, J., Meng, L., et al. (2012). Escherichia coli noncoding RNAs can affect gene expression and physiology of *Caenorhabditis elegans*. Nat. Commun. 3, 1073. doi:10.1038/ncomms2071

Luo, J., He, Q., Xu, J. Z., Xu, C., Han, Y. Z., Gao, H. L., et al. (2021). Microsporidia infection upregulates host energy metabolism but maintains ATP homeostasis. J. Invertebr. Pathol. 107596. Advance online publication. doi: 10.1016/j.jip.2021.107596

Martín-Hernández, R., Bartolomé, C., Chejanovsky, N., Le Conte, Y., Dalmon, A., Dussaubat, C., et al. (2018). *Nosema ceranae* in *Apis mellifera:* a 12 years postdetection perspective. Environ. Microbiol. 20(4), 1302–1329. doi:10.1111/1462-2920.14103

Mayack, C., Natsopoulou, M. E., and McMahon, D. P. (2015). *Nosema ceranae* alters a highly conserved hormonal stress pathway in honey bees. Insect molecular biology, 24(6), 662–670.

McMenamin, A. J., Daughenbaugh, K. F., Parekh, F., Pizzorno, M. C., and Flenniken, M. L. (2018). Honey bee and bumble bee antiviral defense. Viruses. 10(8), 395. doi: 10.3390/v10080395

Naitza, S., and Ligoxygakis, P. (2004). Antimicrobial defences in *Drosophila:* the story so far. Mol. Immunol. 40(12), 887–896. doi: 10.1016/j.molimm.2003.10.008

Paldi, N., Glick, E., Oliva, M., Zilberberg, Y., Aubin, L., Pettis, J., et al. (2010). Effective gene silencing in a microsporidian parasite associated with honey bee (*Apis mellifera*) colony declines. Appl. Environ. Microbiol. 76(17), 5960–5964. doi: 10.1128/AEM.01067-10

Papagiannouli F. (2022). Endocytosis at the crossroad of polarity and signaling regulation: learning from *Drosophila melanogaster* and Beyond. International journal of molecular sciences, 23(9), 4684. doi: https://doi.org/10.3390/ijms23094684

Paris, L., El Alaoui, H., Delbac, F., and Diogon, M. (2018). Effects of the gut parasite *Nosema ceranae* on honey bee physiology and behavior. Curr. Opin. Insect Sci. 26, 149–154. doi: 10.1016/j.cois.2018.02.017

Park, D., Jung, J. W., Choi, B. S., Jayakodi, M., Lee, J., Lim, J., et al. (2015). Uncovering the novel characteristics of Asian honey bee, *Apis cerana*, by whole genome sequencing. BMC Genomics. 16(1), 1. doi: 10.1186/1471-2164-16-1

Pelin, A., Selman, M., Aris-Brosou, S., Farinelli, L., and Corradi, N. (2015). Genome analyses suggest the presence of polyploidy and recent human-driven expansions in eight global populations of the honey bee pathogen *Nosema ceranae*. Environ Microbiol. 17(11), 4443–4458. doi: https://doi.org/10.1111/1462-2920.12883

Pillai, R. S., Bhattacharyya, S. N., Artus, C. G., Zoller, T., Cougot, N., Basyuk, E., et al. (2005). Inhibition of translational initiation by Let-7 microRNA in human cells. Science. 309(5740), 1573–1576. doi: 10.1126/science.1115079

Rabuma T, Gupta OP, Chhokar V. (2022). Recent advances and potential applications of cross-kingdom movement of miRNAs in modulating plant’s disease response. RNA Biol. 19(1): 519–532. doi: 10.1080/15476286.2022.2062172

Robey, R. B., and Hay, N. (2006). Mitochondrial hexokinases, novel mediators of the antiapoptotic effects of growth factors and Akt. Oncogene. 25(34), 4683–4696. doi: 10.1038/sj.onc.1209595

Robinson, M. D., McCarthy, D. J., and Smyth, G. K. (2010). edgeR: a Bioconductor package for differential expression analysis of digital gene expression data. Bioinformatics. 26(1), 139–140. doi: 10.1093/bioinformatics/btp616

Samaranayake, Y. H., Samaranayake, L. P., Pow, E. H., Beena, V. T., & Yeung, K. W. (2001). Antifungal effects of lysozyme and lactoferrin against genetically similar, sequential Candida albicans isolates from a human immunodeficiency virus-infected southern Chinese cohort. J. Clin. Microbiol., 39(9), 3296–3302. doi: 10.1128/JCM.39.9.3296-3302.2001

Schottelius, J., Schmetz, C., Kock, N. P., Schüler, T., Sobottka, I., Fleischer, B. (2000). Presentation by scanning electron microscopy of the life cycle of microsporidia of the genus *Encephalitozoon*. Microbes Infect. 2(12), 1401–1406. doi: 10.1016/s1286-4579(00)01293-4.

Smoot, M. E., Ono, K., Ruscheinski, J., Wang, P. L., and Ideker, T. (2011). Cytoscape 2.8: new features for data integration and network visualization. Bioinformatics. 27(3), 431–432. doi: 10.1093/bioinformatics/btq675

Traver, B. E., and Fell, R. D. (2011). *Nosema ceranae* in drone honey bees (*Apis mellifera*). J. Invertebr. Pathol. 107(3), 234–236. doi: 10.1016/j.jip.2011.05.016

Traver, B. E., and Fell, R. D. (2012). Low natural levels of *Nosema ceranae* in *Apis mellifera* queens. J. Invertebr. Pathol. 110(3), 408–410. doi: https://doi.org/10.1016/j.jip.2012.04.001

Vávra, J. (1976). Structure of the microsporidia. Biology of the microsporidia. Springer, Berlin

Weiberg, A., Wang, M., Lin, F. M., Zhao, H., Zhang, Z., Kaloshian, I., et al. (2013). Fungal small RNAs suppress plant immunity by hijacking host RNA interference pathways. Science. 342(6154), 118–123. doi: 10.1126/science.1239705

Widmann, C., Gibson, S., Jarpe, M. B., and Johnson, G. L. (1999). Mitogen-activated protein kinase: conservation of a three-kinase module from yeast to human. Physiol. Rev. 79(1), 143–180. doi: 10.1152/physrev.1999.79.1.143

Wu, Y., Zheng, Y., Chen, Y., Chen, G., Zheng, H., and Hu, F. (2020). *Apis cerana* gut microbiota contribute to host health though stimulating host immune system and strengthening host resistance to *Nosema ceranae*. R. Soc. Open. Sci. 7(5), 192100. doi: https://doi.org/10.1098/rsos.192100

Xing, W. H., Zhou, D. D., Long, Q., Sun, M. H., Guo, R., Wang, L. M. (2021). Immune response of eastern honey bee worker to *Nosema ceranae* infection revealed by transcriptomic investigation. Insects. 12(8), 728. (in Chinese)

Xiong, C. L., Chen, H. Z., Geng, S. H., Zhou, N. H., Zhou, D. D., Zhu, Z. W., Chen, D. F., et al. (2020). Expression profile of high-expressing genes and its potential role during *Apis cerana cerana* infected by *Nosema ceranae*. Journal of Sichuan University(Natural Science Edition), 57(03), 596–604. (in Chinese)

Xiong, C. L., Du, Y., Feng, R. R., Jiang, H. B., Shi, X. Y., Wang, H. P. et al. (2020). Differential expression pattern and regulation network of microRNAs in *Ascosphaera apis* invading *Apis cerana cerana* 6-day-old larvae, Acta Microbiologica Sinica, 5, 992–1109. (in Chinese)

Yang, D. L., Pan, L. X., Chen, Z. Z., Du, H. H., Luo, B., Luo, J., et al. (2018). The roles of microsporidia spore wall proteins in the spore wall formation and polar tube anchorage to spore wall during development and infection processes. Exp. Parasitol. 187, 93–100. doi: 10.1016/j.exppara.2018.03.007

Zhang, L., Hou, D. X., Chen, X., Li, D. H., Zhu, L. Y., Zhang, Y. J., et al. (2012). Exogenous plant MIR168a specifically targets mammalian LDLRAP1: evidence of cross-kingdom regulation by microRNA. Cell Res. 22(1), 107–126. doi: 10.1038/cr.2011.158

Zhang, T., Zhao, Y. L., Zhao, J. H., Wang, S., Jin, Y., Chen, Z. Q., et al. (2016). Cotton plants export microRNAs to inhibit virulence gene expression in a fungal pathogen. Nat. Plants. 2(10), 16153. doi: 10.1038/nplants.2016.153

